# Curved crease origami and topological singularities at a cellular scale enable hyper-extensibility of *Lacrymaria olor*

**DOI:** 10.1101/2023.08.04.551915

**Authors:** Eliott Flaum, Manu Prakash

## Abstract

Eukaryotic cells undergo dramatic morphological changes during cell division, phagocytosis and motility. Fundamental limits of cellular morphodynamics such as how fast or how much cellular shapes can change without harm to a living cell remain poorly understood. Here we describe hyper-extensibility in the single-celled protist *Lacrymaria olor*, a 40 *µ*m cell which is capable of reversible and repeatable extensions (neck-like protrusions) up to 1500 *µ*m in 30 seconds. We discover that a unique and intricate organization of cortical cytoskeleton and membrane enables these hyper-extensions that can be described as the first cellular scale curved crease origami. Furthermore, we show how these topological singularities including d-cones and twisted domain walls provide a geometrical control mechanism for the deployment of membrane and microtubule sheets as they repeatably spool thousands of time from the cell body. We lastly build physical origami models to understand how these topological singularities provide a mechanism for the cell to control the hyper-extensile deployable structure. This new geometrical motif where a cell employs curved crease origami to perform a physiological function has wide ranging implications in understanding cellular morphodynamics and direct applications in deployable micro-robotics.

**Significance statement:** Here we present the discovery of curved crease origami at the scale of a single cell. We show how topological singularities in the origami (d-cones) and twist walls in microtubule ribbons control deployment of a hyper-extensile neck in a single-celled protist. Our work establishes a direct link between geometry and cell behavior, connecting form and function of cellular morphodynamics.

## 1 Introduction

Unicellular protists are mesmerizing to observe as they perform dramatic morphological changes in real time. Large physical transformations in cell architecture have been well-studied during division [1, 2, 3], or in motility, such as when cells crawl through narrow pores [4, 5, 6]. Many dynamic behaviors of cells within multi-cellular systems can also be attributed to dramatic contractile and extensile properties, such as that of cardiac fibroblasts in the heart [7], peristalsis in the gut [8], or the extreme stretchability without rupture of the lung epithelium [9]. Individual cells must undergo large strains and strain rates to accomplish these behaviors [10].

What sets the fundamental limits of cellular morphodynamics remains unknown. Here we use hyper-extension in a predatory ciliate *Lacrymaria olor* [11, 12, 13, 14, 15] to establish a system to study the limits of morphodynamics in single cells. To the amusement of an observer, a small 40 *µ*m single cell is capable of extending a neck-like protrusion up to 1500 *µ*m in less than 30 seconds, then retracting this neck as quickly as it was extended [16, 17, 18, 19, 20, 21]. Our previous work established the role of these extensions in comprehensively searching its surroundings for prey [15, 22]. As opposed to passive cellular deformations [23, 24], *L. olor* actively extends and retracts its neck-like protrusion [15, 22] via ciliary dynamics. It is a remarkable feat of large scale deformations of a single cell that *L. olor* repeats more than 20,000 times in its lifetime without failure. While cells can have many different types of subcellular protrusions such as cilia, flagella, microvilli, axons, and more [25], the reversible, rapid, hyper-extensible protrusion of *L. olor* is unique, and the underlying mechanisms which produce the cell’s extreme morphing behavior remain unknown. Previous attempts to explain shape change in protists explored the role of shear amongst filaments to modulate Gaussian curvature [26, 27]. Although relevant to Euglena, an extension of 30 times the body length, as we report here in *L. olor*, is not feasible via pure shear alone.

Our current work aims to link form and function in a dynamic shape-shifting protist highlighting the role of geometry in programming unicellular behavior [28]. Piecing together data from multiple imaging modalities including real-time imaging and TEM ultra-structure characterization, we discover a novel “curved crease origami” architecture within ultra long helical microtubule ribbons that support hyper-extensibility. We demonstrate using scaled-up physical models that this unique origami architecture of the cell directly enables ultra-long reversible strains and large strain rates via geometrical transformations driven by ciliary activity. Curved crease origami further creates a readily-deployable mechanism by controlling a sharp transition in cell geometry between folded and unfolded zones using well known d-cone and twisted domain wall singularities. This deeper understanding of how cortical cytoskeleton enables transformable material properties has the potential to expand our capacity to build micro-scale living machines and deployable structures [25].

## 2 Results

### 2.1 Comparative Single Cell Extensibility

Long thin protrusions are a common motif in biological cells. The largest strains recorded at the cellular level often occur in thin extensile protrusions [25]. For example, spinal motor neuron axons can grow up to 1 m in humans, compared to their cell body size of 10 µm [29, 30]. Similarly, the highest strain rate recorded in a single cell also occurs in an extensile protrusion in ultra-fast discharges of nematocysts that occur at speeds up to 18.6 m/s [31]. We compiled published data on a variety of extensile protrusion dynamics in single cells (Fig. 1, Supplementary Table 1) to observe the diversity of strain and extension rates across the eukaryotic tree of life. At one extreme are ultra-fast single-shot extensions such as in Microsporidia which use eversion of tube geometry to enable fast extrusion of tube-like structures [32, 33, 34]. Conversely, several cell types such as neuronal cells are able to extend to great lengths but only on longer developmental timescales [35]. Additionally, most cellular protrusions are often irreversible [25]. One cell that stands out with both reversible and fast extensile protrusions is *Vorticella*, a single cell filter feeder with a long contractile stalk [36, 37]. Though fast, the reversibility is still limited since extension is orders of magnitude slower than contraction. Additionally, the total extension length is significantly limited compared to other types of extensile protrusions [36]. Comparing this data set to both wild type and lab cultures of the ciliate *L. olor*, we find rapid reversible “neck-like” protrusions that can reach lengths 30 times the size of the cell (40 *µ*m) at extension and contraction speeds of 1000 *µ*m/s. This is a remarkable reversible extension of a single cell and immediately raises many questions about how a cell can generate that much membrane and cortical cytoskeleton to support such a large extension. Our current work directly addresses this question.

**Figure 1:**
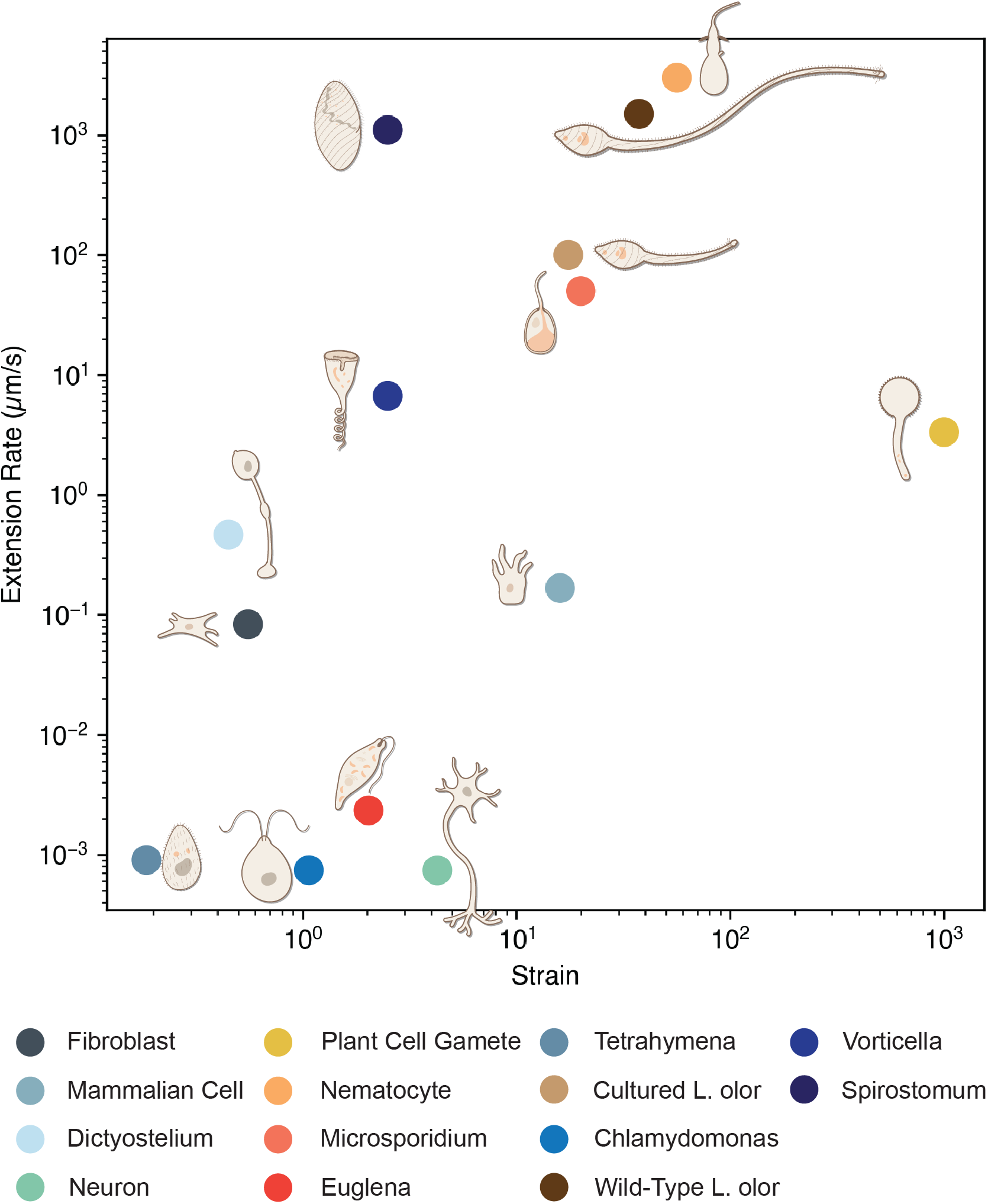
Hyper extensible single cells with large strain and strain rates. Extension rate (µm/s) versus strain (ΔL/L) of protrusions in both unicellular and multicellular cell types. All cell types, and the name of their extensible protrusions when applicable from bottom to top: *Chlamydomonas* (cilia)[90], *Tetrahymena* (cilia)[90], *Euglena*[91], Neuron (axon)[35], Fibroblast (lamellipodium)[92], Sea Urchin Embryonic Cell (filopodium)[93], *Dictyostelium* (pseudopod)[94], Maize Gamete (pollen tube)[95], *Vorticella* (stalk)[37], Cultured *Lacrymaria olor* [15], *Spirostomum*[96], Wild-type *Lacrymaria olor* [15, 38, 39], Nematocyte (nematocyst)[33], *Microsporidium* (polar tube)[32].

### 2.2 Hyper-extensibility in a single cell

*Lacrymaria olor* is a unicellular ciliate in the family Lacrymariidae. This predatory ciliate presents a unique morphology with a bulbous body often attached to substrates and an active neck-like protrusion that searches for prey. As the neck extends and retracts, cytoplasm containing small granules can be seen rushing in and out of the neck while a majority of the cytoplasm and the cell nucleus remain anchored in the main cell body [15]. The body and neck are uniformly covered with cilia which enable motility and neck length dynamics [22, 15]. Additionally, the oral apparatus at the anterior “head” contains specialized longer cilia which collectively beat or pause regularly. In case of encounter with prey, the neck behavior changes dramatically leading to remodelling of the neck [38, 39]. Formation of a food cavity in the neck once prey is captured leads to a bulge which is supported by the cortical cytoskeleton as the prey is reeled in towards the cell body [38, 39, 15].

Previously, we reported on analysis of neck extension/contraction dynamics in lab cultures [15]. Surprisingly, wild collected cells show hyper-extension on a regular basis. During the search for prey, the neck-like protrusion in wild-type cells demonstrates hyper-extension [16, 17, 18, 19, 20, 21]). For example, a single cell is capable of an extension reaching 1210 *µ*m in under 7 seconds while maintaining a constant cell body size of 40*µ*m (Fig. 1, Fig. 2A-C, [16, 17, 18, 19, 20, 21]). *L. olor* can then fully retract this protrusion as quickly as extended and re-extend immediately (Fig. 2A-C, [16, 17, 18, 19, 20, 21]). We have recently linked these extension/contraction dynamics directly to the search cloud observed in *L. olor* [22]. After a burst of extension/contraction activity, the cell neck fully retracts and the cell becomes “dormant” for a few minutes, remaining in this state until its next hunting event [15].

**Figure 2:**
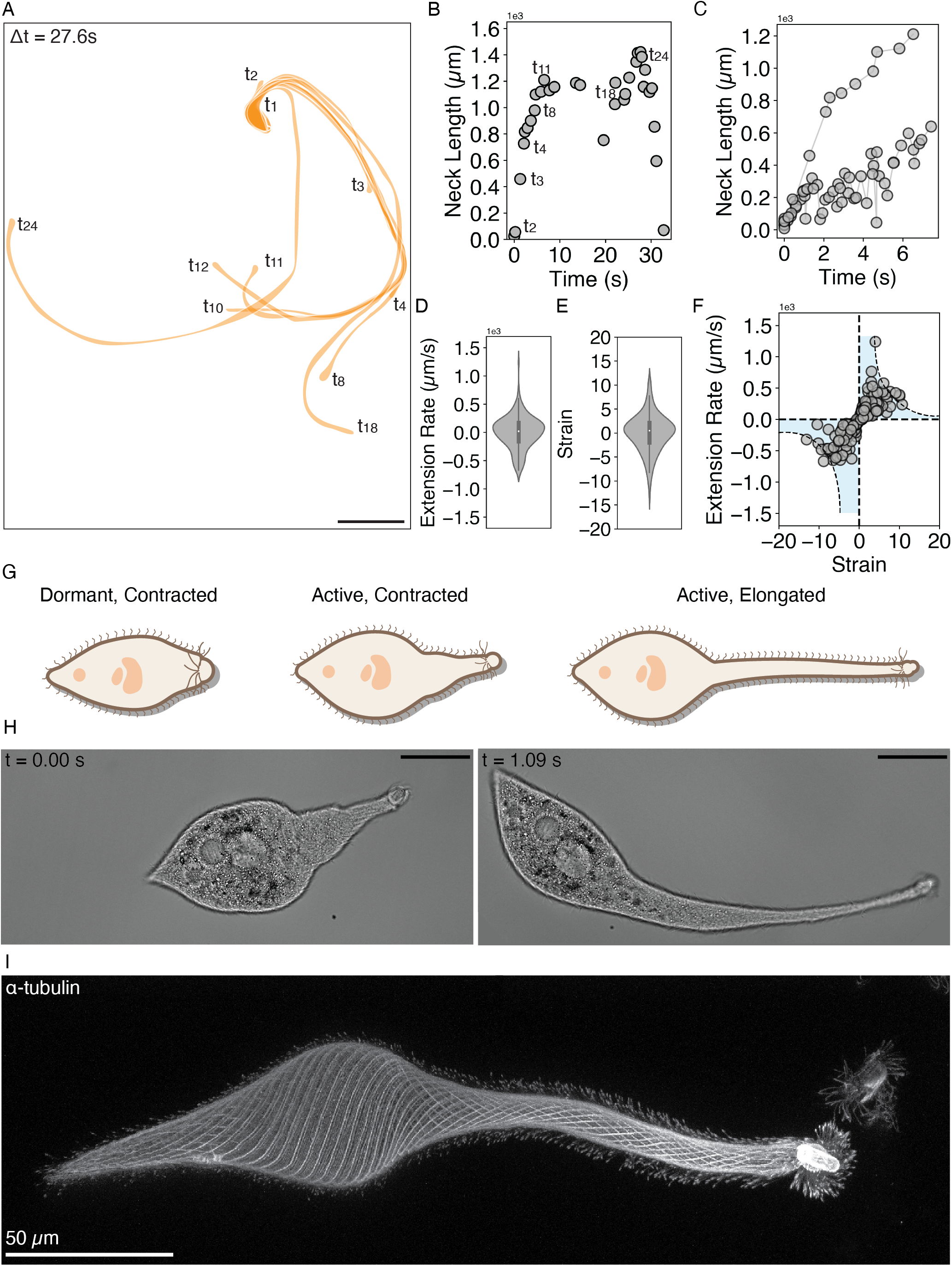
Lacrymaria olor undergoes extreme extensions at ultra fast speeds. (A) An overlay of snapshots from one neck extension-contraction cycle over a duration of 27.6 seconds shows an active cell which can reach many unique points around its cell body. Scale bar is 150 *µ*m.(B) Starting from an active, contracted state, the cell can reach an extension of 1210 *µ*m in under 7 seconds then retract just as quickly. (C) Time-points from multiple extension events during an active hunting period show similar extension rates. (D) Extension [+] and contraction [-] rates calculated from these extension events show an extension rate of up to 1420 µm/s. Individual extension/retraction events are noisy while the overall maximum extension rates are slightly faster than retraction rates. (E) Strain calculated for these deformations during extension [+] and contraction [-] show large strain magnitudes [15+] for a single cell. (F) Extension Rate vs. Strain for these same extension events. The blue shading and dotted lines were drawn in to highlight the reversibility of these dynamics. Data in A-F analyzed from publicly available wild type cell videos accessible at [16, 17, 18, 19, 20, 21]). (G) *L. olor* transitions between a dormant, contracted state and two active behavioral states by extending and contracting a neck-like protrusion. (H) The distinct cell morphologies between active contracted and active elongated states are apparent in live cells under DIC imaging. Scale bar is 40 *µ*m. (I) Confocal fluorescence z-stack projection of *α*-tubulin stained fixed cells characterizes the helically-arranged cortical cytoskeleton.

Tracking cell shape as a function of time (Fig. 2A-C, [16, 17, 18, 19, 20, 21]) reveals fast extension dynamics (1200 microns in 7 seconds) with almost equally fast retraction speeds (Fig. 2D-F). Such a fast, reversible extension has never been reported for any other living material, let alone a single cell. This raises the question of how a single cell can achieve such extension and retraction rates.

### 2.3 Cortical cytoskeleton as a base for non-affine cellular exten-sions

To understand the mechanics underlying this ultra-fast extension and contraction, we utilized high speed DIC imaging to directly observe the neck extension process in live organisms (Phantom v1210, 200fps, see methods for details) (Fig. 2H, Supplementary Movie 1). When undergoing the transition from a dormant state to an active hunting state, the cell elongates to its longest neck lengths through a piece-wise process with pauses coinciding with ciliary reversals (Fig. 2G,H, Supplementary Movie 1). The cell’s initial extension from dormancy is a slow extension into a transition state, where the neck is only partially elongated (Fig. 2G). Once in this state, the neck can then elongate rapidly, and the cell can easily traverse morphologically between an active contracted state and an active elongated state. It is important to note that although a dramatic extension of the cell has occurred during this transition, much of the main cell body is hardly transformed. Thus this neck extension is not a classical Hookean extension, but instead a non-affine extension where the neck elongates without any extension observed in the cell body.

Since ciliates are defined classically by their cortical cytoskeleton structures which also provide a map for surface cilia [40], we next mapped the cortical cytoskeleton structure of *L. olor*. During these extensions captured using high speed DIC imaging, the underlying cortical cytoskeleton is also visible (Fig. 2H). The neck and cell body of *L. olor* are structurally supported by a cortical cytoskeleton primarily consisting of microtubule bundles (Fig. 2I) and centrin (Fig. 3H) [14, 15]. Microtubule filaments in an extended cell weave a left handed helical structure, forming continuous bands from the tail of the cell body to the apical tip of the neck (Fig. 2I, 3A). The presence of filaments even in long necks and their continuation into the body of the cell provides the first hint that the cortical cytoskeleton might play a role in enabling long neck extensions.

**Figure 3:**
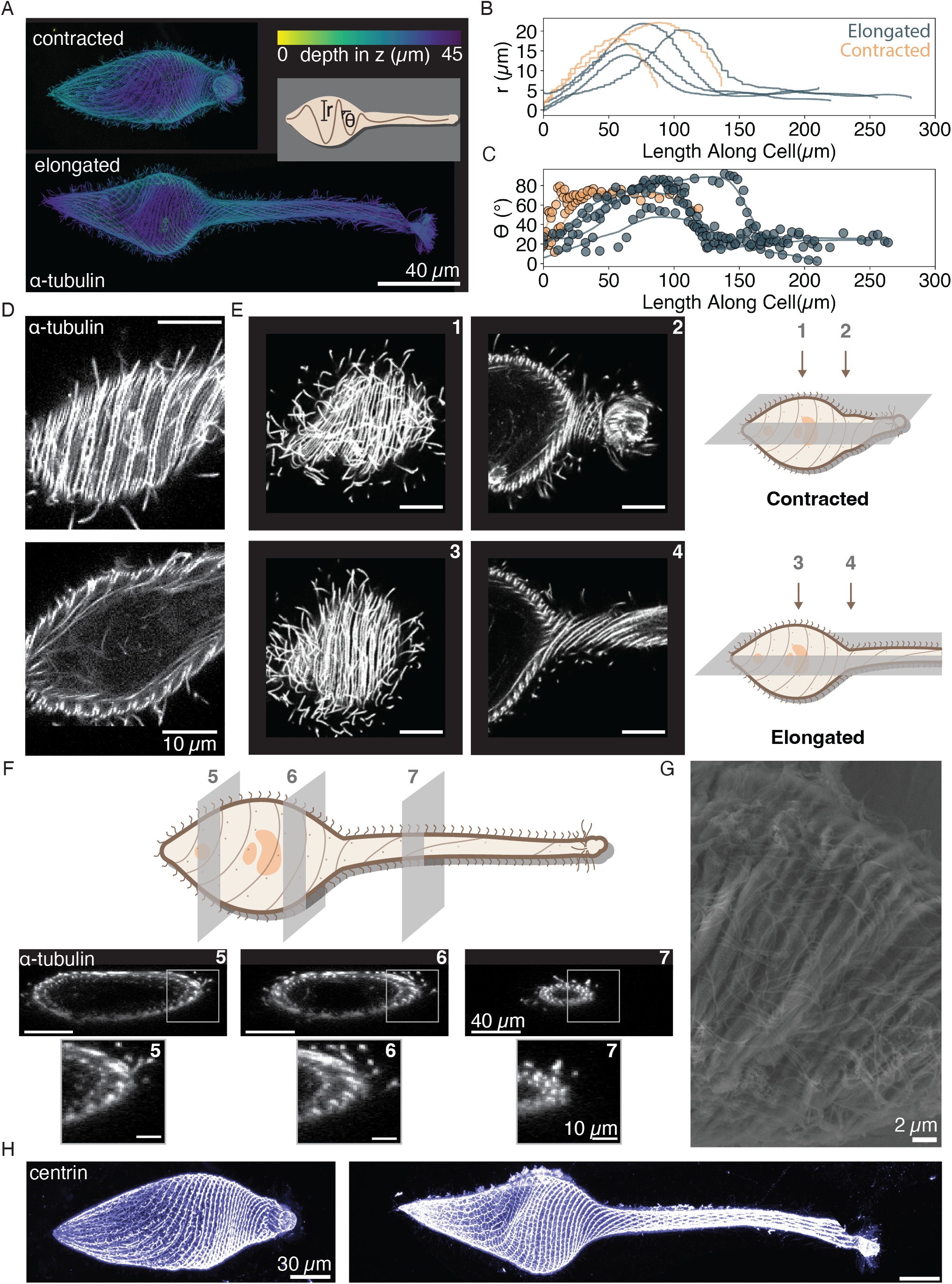
A helically-arranged cortical cytoskeleton contains layers of microtubule filaments amid membrane folds. (A) Confocal fluorescence z-stack projections of *α*-tubulin in fixed contracted and elongated cells. Inset shows measurements taken on these 2d projections of z-stack projections (radius and helix angle).(B) The microtubule structure varies in radius along the length of the cell in both elongated and contracted cells (n=6). (C) The microtubule structure varies in helix angle along the length of the cell in both elongated and contracted cells (N=6). (D) Microtubule filaments occur in doublets which are additionally layered in z. Scale bars are 10*µ*m. (E) These layers are present both in the body and neck region in contracted cells, and in the body region of elongated cells, however they are not observed in the neck region of elongated cells. Scale bars are 10*µ*m. (F) Schematic showing the planes in panels 5-7, which are three rotated images of the *α*-tubulin in three different planes along the length of the cell. Panels 5-7 confirm multiple layers of *α*-tubulin in the cortical cytoskeleton. Below are zoom-ins of the images which highlight the layers. (G) Scanning Electron image of the body of an elongated *L. olor* cell. (H) Confocal fluorescence z-stack projection of centrin stained fixed cells. Scale bars are 30 *µ*m.

Next we label cortical cytoskeleton of a broader range of ciliates with similar shape as *L. olor*. We find that other single-celled ciliates with similar cellular morphologies have dramatically different cortical cytoskeleton architectures and dramatically different cell behaviors. *Dileptus*, for example, has a tear-drop shaped cell body and a neck-like protrusion which the cell is not able to elongate or contract. The cortical cytoskeleton primarily consisting of microtubule filaments is aligned along the long axis of the cell (Fig. S1). *Tracheloraphis*, on the other hand, does not have a neck-like protrusion, but does exhibit whole cell contraction and elongation. This cell similarly has a cortical cytoskeleton composed of microtubule filaments which again align with the long axis of the cell (Fig. S1). The different architectures and behaviors in morphologically similar cells further supports our initial hypothesis that the cortical cytoskeleton architecture in *L. olor* plays a role in the rapid, reversible hyper-extension observed.

We next search for what allows microtubule filaments to be present in the extended neck of *L. olor*. Although microtubule filaments are known to undergo dynamic instability, the growth and catastrophe rates (10 *µ*m/min) of those microtubule polymerization processes are too slow to account for the 1000 *µ*m/s neck elongation in *L. olor* [41, 15]. If not polymerized and depolymerized during rapid extension and contraction cycles, pre-existant microtubule filaments must be deployed and recovered for extension events. For an extension similar to the one in Figure 2A, this would require storage up to 1400*µ*m of microtubule filaments within the cell. To understand the upper bounds of this as a possibility, we estimated the total amount of microtubules which could be wrapped around a cylinder whose dimensions are roughly equivalent to a contracted *L. olor* (Supplementary Methods, Fig. S5). Assuming a maximum surface packing of a helical rod, we found that a single layer spool of microtubule filament is capable of storing over 100 times more microtubule length than the maximum neck length observed (see SI for details). This simple calculation points towards potential helical spooling as a mechanism for cytoskeleton storage.

In addition to cortical cytoskeleton, the rapid elongation-retraction cycles also require cell surface area to increase rapidly. Since increasing the surface area of the cell through membrane synthesis occurs too slowly [42] to account for real time neck elongation and contraction in *L. olor*, extra membrane must be present in the cell body prior to extension. The most likely possibility is storage of extra membrane via membrane folding [43, 44, 45, 46]. To approximate how much membrane would need to be stored when in the contracted state, we estimated the surface area of a 1400 *µ*m neck to be 22,000 *µ*m^2^ (Supplementary Methods). However, the surface area of the cell body with a 40 *µ*m diameter has a much smaller surface area of about 5000 *µ*m^2^. This would indicate that roughly 22,000 *µ*m^2^ or at least four-fold extra membrane would need to be stored in the cell when the neck is fully contracted in our current membrane-storage hypothesis.

### 2.4 Multi-layered storage of microtubule bundles in the cortical cytoskeleton

To learn how the microtubules might be stored within the cell, we observed the cortical cytoskeleton in *L. olor* using confocal fluorescence microscopy (Fig. 3). While all ciliates have a cortical cytoskeleton with low-angle helices (i.e. Paramecium, Stentor, Dileptus, Spirostomum) [47], *L. olor* has a a unique helical cortical cytoskeleton architecture which varies in helix angle and radius both along the length of the cell and across contracted and elongated states (Fig. 3A-C,Fig. S2) [14, 15]. By tracing filaments from experimental data (see methods), we directly measure this helix angle which increases with the cell diameter from 20° at the tail up to 60° at the cell body (Fig. 3B,C). In elongated cells, the helix angle again increases starting at the tail up to 80° and then drops rapidly to 20° in the neck continuing through to the oral apparatus (Fig. 3B,C). This decrease in helix angle at the base of the neck similarly coincides with the decrease in axi-symmetric diameter of the cell from 40 *µ*m to 10 *µ*m (Fig. 3B,C). How microtubules are able attain such high helix angles remains an open question given their large persistence length [48] due to high stiffness. Potentially, the source of bending in *L. olor* is assisted by microtubule associated proteins similar to those in microtubule bundles or cilia [49, 50].

We term the zone of dramatic change in cell morphology and cytoskeleton as the “transition zone” of the cell, where the Gaussian curvature of the cell transitions from positive over cell body to negative during transition and zero over the cylindrical neck. As the neck elongates, the microtubule filaments which transition from the body to the neck undergo a large geometrical transformation from high-angle helicity and a large helix radius to low-angle helicity and a small helix radius. For the rest of the work, we focus specifically on this transformation and the “transition zone” as both membrane and microtubule network change confirmation via this zone.

Higher resolution three dimensional imaging further reveals that what appeared to be a single filament at low magnification is actually a pair of filaments, and that these pairs of filaments are present in multiple layers (spools, Fig. 3D). This layering was most apparent in the cell body (Fig. 3E-F), and is no longer present in the neck of elongated cells (Fig. 3E). We confirmed this observation by rotating confocal z-stack data-sets to look head on into the cell. In both the body and the transition zone from the body to the neck two layers of filament pairs were visible (Fig. 3F, Supplementary Movie 2). The lack of microtubule layering in the neck of an elongated cell, and the presence in the body and transition zone of both cells (Fig. 3E,F) suggests that the layering is likely a mechanism of microtubule spooling within the cell. We also observed that the microtubule bundles pair together with ciliary pits lining between them (Fig. 3D-E), anchoring ciliary basal bodies to the helical architecture. Thus the pitch and hence the cilia beat force change dramatically as the cytoskeleton geometry evolves with an extending neck. For a higher helical pitch, much of the ciliary beat is aligned along the radial axis while lower helical pitch aligns ciliary beats along the long axis of the cell. This geometrical feedback between cell architecture and surface activity couples hydrodynamic forces generated by individual cilia in the direction of neck extension. Thus the number of cilia involved in neck extension increases and scales with neck length.

To the best of our knowledge multi-layer spooling of microtubules has never been reported in a cellular context before. A simple three layer spooling of microtubules can easily triple the total length that can be stored in a cell body (See SI for details), allowing the microtubules to be ready for deployment during extensions. The presence of multi-layered microtubules removes the need for filament growth, allowing the cell to store a sufficient length of microtubules filaments to account for the longest neck length extensions observed (See SI for details, Supplementary Movie 2).

Furthermore, since centrin, a calcium-binding protein, is often found in the cortical cytoskeleton of many ciliates [51, 52, 53], we further looked for centrin distribution in the cortical cytoskeleton of *L. olor*. As shown in Figure 3H, centrin forms a highly networked cross-linked (mesh-like) structure following the same distribution as microtubules described above (Fig. 3H). The driving force for neck contraction has been hypothesized to be centrin [15, 22] which is known to have a calcium-dependent contractile phenotype in a mesh-like form [52, 53, 51]. Beyond it’s potential role in contraction processes for the neck, such a linked cortical cytoskeleton is conducive to inelastic and inextensible sheet-like behavior by allowing for both in-plane rigidity and out-of-plane bending stiffness. As shown in the imaging datasets, the centrin architecture deforms due to extension but is not disrupted by repeated cyclic extension and contraction of the neck (Fig. 3H). Inextensibility of the membrane in this context will become relevant in the next few sections.

### 2.5 Membrane storage in pleats at the transition zone

To understand how the membrane might be stored and redeployed within the cell, we initially performed Scanning Electron Microscopy (SEM) on a fixed cell in an extended state and observed the presence of membrane folds in the body of the organism. These folds correlate exactly to the previously mapped cortical cytoskeleton (Fig. 3G). Next, we performed fluorescence imaging using FM4-64 (a membrane marker) on fixed cells and found that the membrane folds follow a similar helical trajectory as we described previously for microtubules (Fig. S3).

To further quantify the degree of membrane storage and geometry of these membrane folds, next we performed Transmission Electron Microscopy (TEM) on fixed cells (see methods section for details). We found clearly defined membrane pleats as opposed to disorganized ruffles seen in many eukaryotic cells (Fig. 4A-B)[43, 54]. The pleats are uniform and periodic in the contracted cell, with the depth of pleats ranging from 3*µ*m in the tail to 2 *µ*m in the transition zone (Fig. 4C). Remarkably, we also saw a clear transition from a pleated to flattened geometry roughly the depth of 1 *µ*m exactly at the “transition zone” in an elongated cell (Fig. 4B,D). 1 *µ*m is also the minimal depth of the ciliary pits in the organism (Fig. 4C-D).

**Figure 4:**
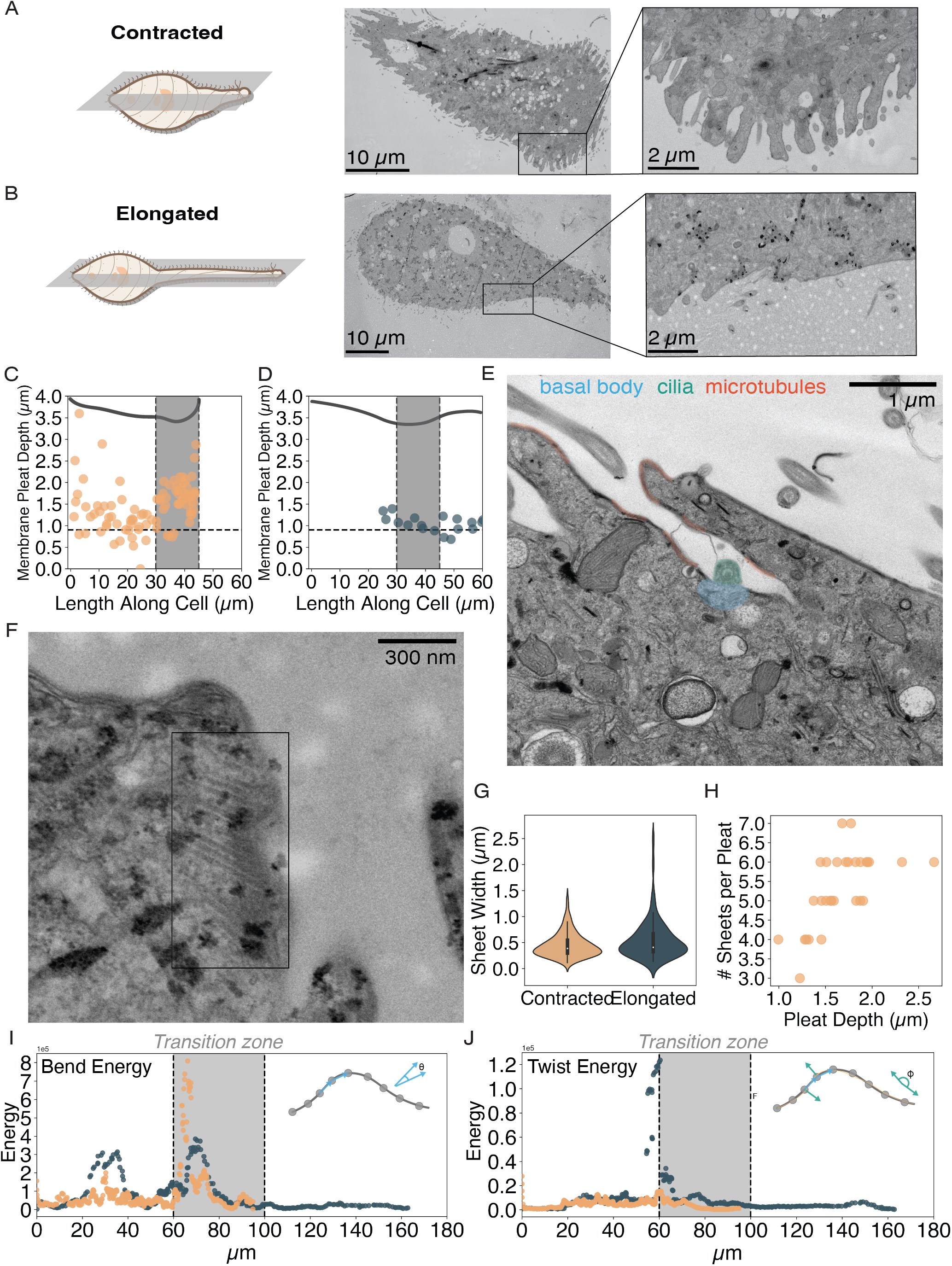
The membrane is folded into pleats which contain contain microtubule ribbons. (A-B) TEM images of slices in fixed contracted and elongated cells highlight the presence of membrane folds in the body-to-neck region of a contracted cell which are absent in the same region of an elongated cell. (C) The pleat depth in a contracted cell increases in the body-to-neck region (transition zone). The transition zone is highlighted in grey, and the region of the transition zone within the cell is shown using an outline of the cell in the top of the plot. (D) The pleat depth remains the minimal depth of the ciliary pit depths in the same zone in an elongated cell. The transition zone is highlighted in grey, and the region of the transition zone within the cell is shown using an outline of the cell in the top of the plot. (E) Representative TEM image of a membrane pleat show that these folds contain ciliary pits. (F) There are many microtubule ribbon in each of these pleats which lie along the membrane. (G) The median width of a ribbon is consistent in both contracted and elongated cells. (H) The total number of ribbons per pleat increases with pleat depth in contracted cells. (I-J) Microtubule bending and twisting creates an energy barrier at transition zone.(I) Schematic shows how the bending energy was calculated from the points in each filament. The bending energy has a global maximum in the transition zone in both a contracted and an elongated cell, with a higher energy maxima in the contracted cell. (J) Schematic shows how the twisting energy was calculated from the points in each filament. The twisting energy has a global maximum at the beginning of the transition zone an elongated cell, with a lower energy maxima in the contracted cell.

The structure and localization of these membrane pleats allow us to create a more quantitative definition of the transition zone. We had previously defined the transition zone as the region of slight neck elongation in the active, contracted state of the cell. This new data allows us to define the transition zone as the region where the membrane transitions from pleated (folded) to straight (unfolded) in an elongated cell, which is highlighted in gray in Figures 4C-D. This transition from folded to unfolded membrane pleats rapidly provides the much needed membrane for long extensions in real-time (See Supplementary Movie 1).

### 2.6 Integrated ultrastructure of the membrane and the cortical cytoskeleton

Looking more closely at the TEM sections, we additionally observe that the membrane pleats harbor individual cilia (Fig. 4E). The basal bodies are at the bottom of these pleats, with cilia visibly extending from them (Fig. 4E). Additionally, we observed bundles of microtubule filaments lining these membrane pleats (Fig. 4E, highlighted in orange). A closer look at one of these bundles in Figure 4F shows that these bundles are each actually a ribbon of microtubule filaments, and they are localized and anchored with the membrane (Fig. 4F). In both contracted and elongated cells the median width of these microtubule ribbons is 386 nm and 408 nm, respectively. Since these are projections of 3D microtubule helices wound around axi-symmetric structures, TEM cross-sections depict projections that show variation which is an artifact of the sectioning. Knowing the diameter of a single microtubule filament [55], we estimate that the ribbons roughly contain 14 microtubule filaments (Fig. 4G). We further noticed that the microtubule ribbons remain in the same tangential plane of the membrane at all times, providing evidence that they are structurally anchored into the membrane, also seen in other cortical cytoskeleton structures for well studied ciliates [40, 56]. The cortical cytoskeleton is normally also linked to the membrane through basal body linkages which mediate translation of force generation from the ciliary activity to the cortical cytoskeleton [56, 57] (Fig. 4D-E).

The localization of the microtubule ribbons inside the membrane pleats further explains the prior observation of multi-layered microtubules. The association with membrane pleats allows for the microtubule ribbons to spool in a layered architecture and unspool when the pleats unfold in the transition zone as the neck extends. This is further supported by the observation that in contracted cells, the number of microtubule ribbons observed in a membrane pleat decreases as the pleat depth decreases in the neck region (Fig. 4H). Correlating structures we observe both in TEM and confocal data, we identify 15 contiguously wrapped pleats which contain 2-6 microtubule ribbons (depending on the depth of the pleat) of 14 microtubule filaments.

Carefully looking at the pleat structure of the membrane across the whole cell, the architecture can be mapped to an axi-symmetric curved crease origami. This unique curved, pleated membrane architecture and its associated guiding microtubule ribbons simultaneously resolve the issue of membrane and cortical cytoskeleton storage necessary for real time “deployment” of a neck-like extension. This integrated membrane/cytoskeleton architecture is unique and to the best of our knowledge has never been seen before. By establishing this static architecture both in contracted and extended states, we next explore how this transition can occur as a function of neck extension.

### 2.7 Transition zones as energy barriers for neck deployment

We now have a geometrical view of how both membrane and microtubules are stored within the cell enabling large extensions and retractions in short periods of time. However, since microtubule filaments, ribbons and bundles are also known to have large bending and twist energies [48, 58, 59, 60], we anticipate that these tightly curved folds of microtubule ribbons should have energetic consequences. To better understand these potential consequences, we began by characterizing the stored energy in a microtubule ribbon (as described Fig 2,3,4) arranged in the helical geometry of *L. olor* (Fig. S4). We used the exact curves of the microtubule filaments from 3D confocal fluorescence data. We input these curves to compute both the bending and twisting energies of the microtubule ribbons using discrete forms of bend energy 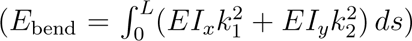 and twist energy 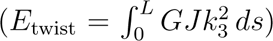 (Fig. 4I,J, Fig. S4, see supplementary methods for details). Interestingly, we observed a peak in locally stored energy right at the transition zone both in bending and twisting energies of filament ribbons in elongated cells, with a second smaller peak in the bend energy near the base of the cell where the radius of the cell decreases rapidly (Fig. 4I-J). Furthermore, in contracted cells, there is a larger peak in bend energy in the transition zone, yet almost no peak in the twist energy (Fig. 4I-J).

This calculation, purely based on microtubule geometry, reveals that an energy barrier exists that the microtubule ribbons must pass through as they transition from the spooled (stored state in a contracted cell) to extended state in an elongated cell via the transition zone. We now can also view this zone as the site of an energy barrier which the microtubule ribbons must pass through. Similarly, the same energy barrier needs to be crossed for the unspooled microtubules to be spooled again when the cell contracts. This geometrical feature at the point of transition zone thus creates a control anchor point which spatially separates the spooled and unspooled states of the elastic (membrane-microtubule complex) system.

### 2.8 Curved crease origami provides a framework for understanding hyper-extensions at cellular scales

Next we examine the role of the “curved crease” pleated origami structure as a control mechanism for deployment of the neck into hyper-extensile states. Although explored as an art form and inspiration for engineering structures and architecture [61, 62, 63, 64, 65, 66], curved crease origami [63] remains the lesser known cousin of traditional origami, where folds are made along straight lines. Starting with work by Huffman [67], a number of key papers have focused on potential developability of curved crease origami structures [64] in inextensible thin sheets of paper [64, 68, 69, 61, 70]. Additionally, various computational and analytical methods have been established to pleat complex curved surfaces [71]. However, only a few studies include mechanical responses of curved crease structures, focusing primarily on simple geometrical shapes and multi-stable systems [69, 72, 73]. Here we open a new realm in curved crease origami and present a unique hyper-extensile origami architecture at “cellular scale”.

As opposed to wrinkled membranes which are disordered and where fold patterns spontaneously emerge [74], the curved creases in the pleated structure of *L. olor* are specifically prescribed by the pattern of microtubule ribbons. Membrane pleats wrap contiguously around the rotationally symmetric cell body, leading to a helicoidal wrapped curved crease pleated structure, as is drawn in Figure 5A. This occurs since stiff microtubules define a “fold line” where the inextensible membrane can only bend along the microtubule ribbon, creating a pleat dictated by microtubule placements.

**Figure 5:**
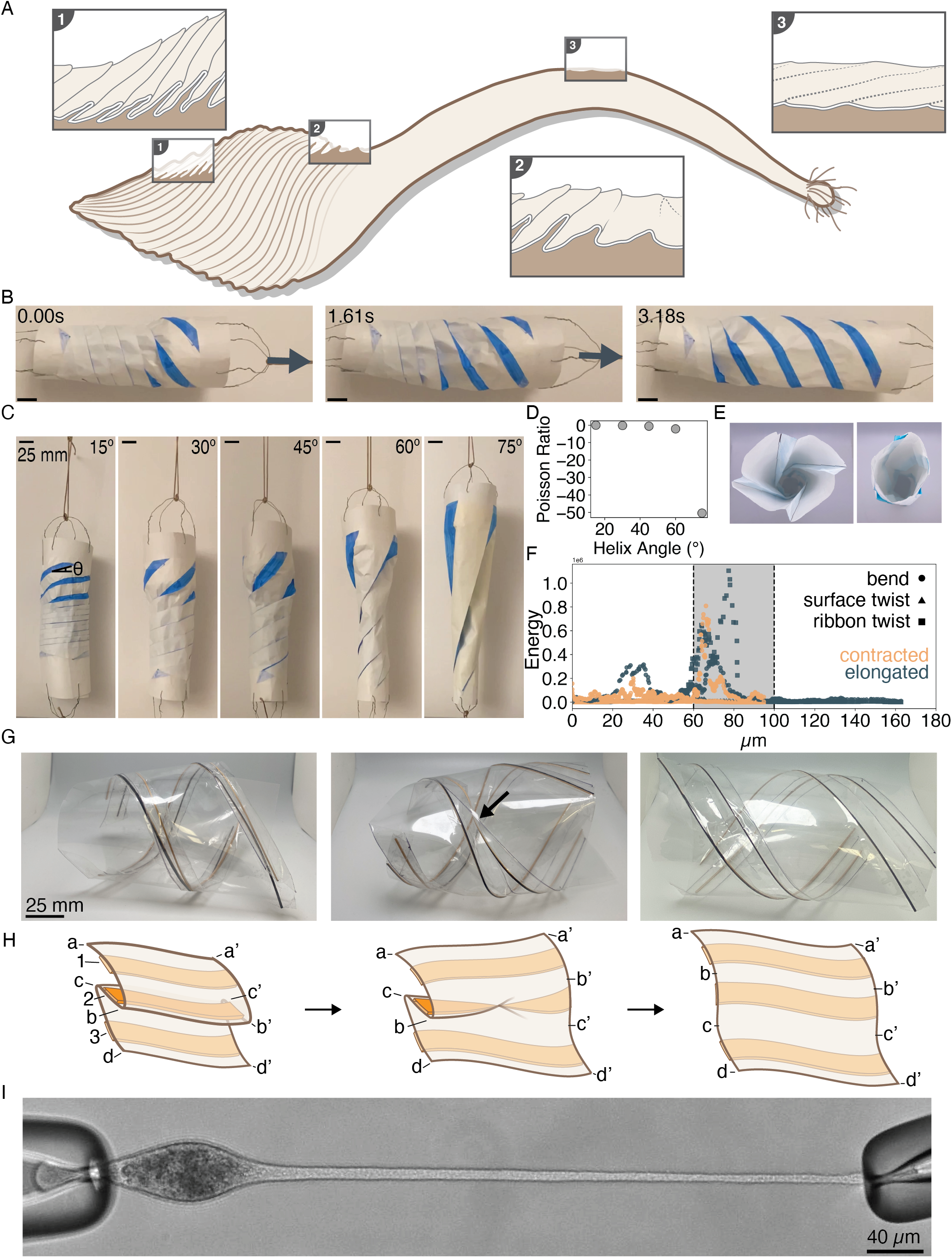
Membrane-microtubule spooling: curved crease origami in a living cell. (A) Schematic of the membrane and microtubule of contracted and elongated cells which highlights the sequential nature of the membrane pleat opening during extension. (1) Zoom-in of the membrane and cortical cy-toskeleton folds in the body of an elongated cell. (2) Zoom-in of the membrane and cortical cytoskeleton folds in the transition zone of an elongated cell. (3) Zoom-in of the membrane and cortical cytoskeleton folds in the neck of an elongated cell. (B) Folded origami structures which are pulled on to elongate. This elongation process demonstrates that the membrane prevents the microtubules from extending like a normal spring through d-cone singularity propagation. The result is sequential opening of the curved crease pleats. Scale bars are 25 mm. (C) Sequential unspooling was observed for multiple fold angles (15*^◦^*,30*^◦^*,45*^◦^*,60*^◦^*,75*^◦^*). (D) The Poisson Ratio for extension at all angle is less than zero. (E) Top-down and Bottom-up views of one origami cylinder show that there is a chirality-induced asymmetry in the pleat structure. (F) Updated energy plot calculated using the geometry of the microtubule filaments in 3D. This updated plot now includes bending energy (circle), twisting energy from the surface of the cell (triangle), and twisting energy from the twisting of the microtubule ribbons (square). (G) Folded cylinders composed of mylar, with bamboo filaments representing the microtubules, and how the ribbons change orientation across folded, transition, and unfolded states. (H) The two d-cone singularities travel, one (B-B’) along a mountain fold and another (C-C’) along a valley fold as force is generated. As the membrane unfolds one of the microtubule ribbons (ribbon 2) twists to accommodate the re-orientation of the membrane. The d-cones and this twist singularity travel as the membrane transitions from folded to unfolded. (I) Image of a cell being held with two micropipettes under suction pressure. The neck was manually elongated while one pipette was kept steady and the second was pulled back. Scale bar is 40*µ*m.

Since origami has not been introduced as a paradigm in a cellular context, we first highlight the unique properties of the combined cortical cytoskeleton and membrane architecture which allow us to model the cell using origami. Traditionally, cell membranes are modeled as thin bilayer sheets which can easily bend but cannot stretch (inextensible) [75]. More importantly, cell membranes also show two-dimensional flow due to in-plane shear on the surface of the cell [76]. In eukaryotic cells, much of this in-plane shear and membrane fluidity is suppressed due to the presence of a cortical cytoskeleton which adheres to the cell membrane [47, 77, 78]. The presence of both microtubule and centrin mesh networks makes the cell membrane in *L.olor* much more like a thin inextensible sheet with very low bending rigidity. This conceptual similarity in properties across thin sheets such as paper and the cytoskeleton-coupled membrane of a cell allows us to explore origami as a conceptual framework to mimic the structure of the membrane folds in *L. olor*.

As the neck elongates, the pleats unfold and smooth out in the neck (Fig. 4) providing the extra membrane needed for real-time deployment of the neck. We schematically depict this unfolding in Figure 5A, where pleats are present in the body of the cell but not in the neck with a sharp change at the transition zone. But how does the cell control where the pleats will open and where they remain folded? In order to better understand this transition, we explore how a helicoidal, pleated, curved crease origami unfolds when stretched.

We created a physical model of membrane storage which mimics that of *L. olor* by scoring a sheet of origami paper at various angles (15°, 30°, 45°, 60°, 75°) (Fig. 5B,C). We then wrapped these sheets into a cylindrical, helical geometry. Here we note that the chirality of the paper origami (seen in Fig. 5B,C, right handed) is opposite of that of the cell (left handed), but that has no impact on the mechanics of this structure. To make clear the stored membrane, we colored the sections between the pleats to represent the membrane that folds and unfolds from the groove. Thus the total colored surface area of the cylinder in this barber-pole-like-origami represents new membrane that is deployed when the cylinder is extended (Fig. 5C, Supplementary Movie 4).

As we unfolded the cylinders, we note that for a single pleat to unfold both the mountain and valley folds have to open (Fig. 5B). Thus, by definition, a single pleat has two associated singularities where the crease goes sharply from folded to unfolded state. These singularities, classically known as d-cones in origami literature [68, 79], travel as the pleat unfolds in the transition zone as a function of time (Fig. 5B). Here we use the word travel in material frame of reference of the cell while the singularity remains stationary in the transition zone in lab frame of reference.

The particular singularity we find at the pleat was first described in context of crumpled paper: a developable cone (d-cone) [79, 80, 81, 68, 82, 83, 84, 85]. This is the simplest of all topological crumpling deformations, where energy focusing introduces a very sharp curvature and much of the strain localizes at the tip. Although the dynamics and interactions of d-cones have been described in literature [84], they have not been studied in the context of pleat folding or unfolding.

Moreover, since these singularities are needed for all pleats to open, each of the 15 pleated grooves along the cross-sectional radius of the cell must have two d-cone singularities in the transition zone. Since d-cone singularities on a surface are coupled by an inextensible material they are known to interact on the surface of a sheet. This unique constraint provides a symmetric neck extension and does not allow one pleat to open without the other 15 pleats also opening synchronously. Any runaway singularity will not be geometrically compatible with the symmetry of the extension, and will slow down to catch up with other d-cone singularities lagging behind. Although it is believed that d-cone singularities coordinate and interact in the classical example of crumpled paper, this coordination of d-cones has not been previously observed [83] in the context of curved crease origami. We directly visualize this process in our high-speed origami unfolding experiments by seeding the initial d-cones and keeping one side of the cylinder (right) open during extension. Because of the nature of the singularity, it is more costly to generate a new singularity at a location further down the helix as compared to just moving the d-cone along the helical path. The singularities in our experiment travel helically in a synchronized manner as the serial unfolding takes place, controlling the extension process (Fig. 5C, Supplementary Movie 4, high speed). Since d-cone propagation requires a critical threshold force [84], this geometrical feature further adds to the energy barrier that controls the deployment of microtubules in the neck. Thus for spooling or deployment to occur, the critical threshold force of extension must be higher than the critical propagation force for all the coupled d-cones.

It is useful to contrast this helical barber-pole-like curved crease origami structure with another classical curved crease pleated origami on a cylinder: a flexible drinking straws [86], where the pleats are concentric rings transverse to the cylinder. These pleats can also store extra membrane and deploy a linear structure while being pulled along the axis of symmetry. However, distinct from the “barber pole curved crease origami” in *L. olor* where singularities control sequential deployment at the transition zone, the pleats here open stochastically and “pop” spontaneously anywhere along the cylinder as the structure is pulled via extension (Fig. S6). This comparison highlights the role of helicoidal geometry in a pleated setting where geometrical singularities control sequential deployment of the spooled membrane and microtubule network during neck deployment.

We additionally observed that there is a chirality induced asymmetry in the cylinders (Fig. 5E). When unconstrained, the helical curved crease origami also shows a unique asymmetry based on chirality of the fold pattern. When the target origami cylinders are observed from the front or from the back end, we observed that there is an asymmetry associated with the ends (Fig. 5E). However, we do not have a mechanistic understanding yet of how this asymmetry emerges.

### 2.9 Twist singularity in microtubule ribbons at the transition zone

To further incorporate elastic structures in the context of curved crease origami of membrane pleats seen in *L. olor*, we next explore the unique role of microtubule ribbons that follow and template the membrane pleats along the grooves. As shown in Figure 4, with each pleat we observed in both TEM and confocal data there were microtubule ribbons on either side of the basal body, which would be on both sides of the fold line in our curved crease origami. Since microtubules are significantly stiffer compared to the membrane (1.2 GPa Young Modulus of microtubules, and by convention a membrane bending rigidity of 20 *k_b_T*)[48], microtubules seed and guide the pleat formation. D-cones moving along the helix can not cross these guiding microtubule tracks.

However, careful observation of this unique ribbon-sheet geometry reveals another coupled singularity during transition of a pleat from folded to unfolded state. Microtubule ribbons present on the inner side of the folded pleat undergo a unique transformation when a d-cone traverses the geometry to unfold the pleats. In order for the pleat to open, one set of microtubule ribbons on the inner pleat must twist 180 degrees, flipping it’s orientation in order to maintain the same face of attachment towards the membrane (Fig. 5H). To depict this geometrical transformation, we made “barber pole” curved crease origami out of transparent mylar and embedded elastic bamboo filaments across the pleat (Fig. 5H). As the cylinder transitions from a folded to an unfolded state, we observed the twist from the uncolored to the colored side of one of the filaments as the d-cones traveled down the curved pleat (Fig. 5G). The twist in the bamboo filaments creates a very large energy barrier, resisting this reorientation due to a twist domain wall singularity. We had previously identified a twist energy barrier of these microtubule ribbons purely from the congruent geometry of the membrane and microtubules (Fig. 4I-J). The discovery of this domain wall is an additional 180 degree twist (occuring in the region bounded by the two d-cones) that increases the contribution of twist energy barrier 10-fold (Fig. 5F, ribbon twist) in the transition zone.

Previously, twist in cytoskeletal filaments has been explored in the context of membrane adhesion energy [87, 88]. This work in bacterial cells led to discovery of unique couplings between the cytoskeleton, membrane curvature, and geometry [87, 88]. In our work, we find that the microtubule ribbon remains attached to the membrane and hence has to twist as pleats open. The above described twist is bounded by the two d-cones which define pleat opening, as shown in Fig. 5H (1,2,3). The creases along which the d-cones travel as the pleat unfolds from left to right are labeled as B-B’ and C-C’. Ribbon 2 undergoes twist, while ribbons 1 and 3 remain in the same plane. The strength of the coupled d-cones defines the length scale of the opening pleats and hence the rate of twist in the microtubule ribbon. This coupling further ensures that this geometry propagates backwards in the material frame of reference of the cell, while remaining stationary in the lab frame of reference (at the transition zone). The constancy of this region in the lab frame during deployment of our physical models also reflects a quasi-static force balance. The cilia-driven pull force is balanced by the energy dissipation of the coupled d-cones and twisting ribbons. Microtubule and membrane pass through the transition zone synchronously while unspooling during deployment (and spooling during recovery), the rate of which is controlled by the singularities and the twisted ribbon. It is important to note, unlike a symmetrical propagation of a d-cone [84], our unique geometry forces d-cones to travel and the ribbons to twist along curved creases, a geometrical case that is yet to be explored. We will further analyze this unique coupled membrane-ribbon geometry and its associated physical properties in a future publication.

### 2.10 Poisson ratio of a deployable curved crease origami

As we observed the extension of cylinders with varying helix angle *θ*, we noticed that in all of the origami experiments the diameter of the cylinder increased post-extension, giving the extensions a negative poisson ratio (Fig. 5C,D). However, during elongation, the neck decreases its net diameter. This can be explained by adding limited shear that a cortical cytoskeletal decorated membrane can support. The models for origami experiment are made of a thin sheets of paper which are incompressible and do not support shear. A thin membrane with bound centrin and microtubules as in *L. olor* is incompressible in 2d as well, but does support limited shear which is driven by ciliary activity. Evidence of this shear extension is present in our cell manipulation experiment.

In order to demonstrate this, we hold a live cell under micro-pipettes both at the neck (oral apparatus) and its tail. Next, we gently extend this cell systematically by application of extension force (see methods). During the neck extension of 47.45 *µ*m, from 126.23 to 173.68 *µ*m in 4.063s, the cell rotates 128.28*^◦^*, rotating at a speed of 31.57*^◦^*/s (Fig. 5I, Supplementary Movie 3), also confirming the chiral nature of the microtubule structure.

Thus hyper-extensibility in *L. olor* is primarily driven by spooling and unspooling of microtubule and membrane architecture along the long axis. A secondary shear provides further yet limited extension along the pull axis. Initially, d-cone propagation generates sequential deployment of microtubule and membrane through the transition zone and into the neck. The d-cone and microtubule unspooling from the pleats give more surface area (membrane storage) and microtubule cytoskeleton during extension. The associated singularities from folded to unfolded states control an organized deployment of the membrane and microtubule network. Initially, this leads to a negative poisson ratio in the cylinder. As the neck is pulled further, a limited shear-induced extension of the neck. During this final extension the neck decreases in diameter, achieving the positive poisson ratio we have observed in *L. olor*.

## 3 Discussion and Outlook

In our work, we share a new mechanism of cellular morphological dynamics which are based on the storage, rapid deployment, and recovery of a neck-like protrusion, and which are harnessed by a cell to create a deployable neck-like structure utilized in capturing prey. In order to understand this hyper-extension in a single cell, we demonstrated a new curved crease origami architecture at the cellular scale. We found 15 curved pleats present in the cell, always folding in a left-handed helix. The pleats connect to the oral grove at the tip of the neck, seeding the d-cone defects that move along the helix during deployment. We mark the key features of our model, including a coupled d-cone and twist singularity that defines the transition zone shown in Fig. 6. Repeatable extension and contraction of this hyper-extensile neck is controlled by the exact synchronized singularities at the transition zone. This property creates a non-affine extensible structure which deploys linearly from the transition zone instead of extending like a simple spring. This further preserves the overall shape of the cell and protects the sub-cellular components in the cell body. The mechanism we describe is reversible, thus the unspooled membrane and microtubule filaments can be spooled back, over and over again. The shapes of many protists are highly dynamic and require fast morphological changes. The principles identified in *L. olor* which enable its unique rapid, reversibile, hyper-extensions can also help us understand the behaviors of other cells or biological systems. Microtubules, for example, are found in the cortical cytoskeleton of all ciliates [89, 47], and the membranes can adopt complex structures which may aid in cellular morphology. Unique centrin architectures have previously been described in other ciliates [51, 52, 53], further adding geometrical constraints on the membrane. Combinations of other types of filaments such as actin with membranous thin sheets can further expand the repertoire of shape dynamics in protists, a rarely studied genre of cells. As high resolution volumetric imaging for a broad range of cells continues to grow, our approach for linking cytoskeletal and membrane geometry to cell function will be broadly applicable.

**Figure 6:**
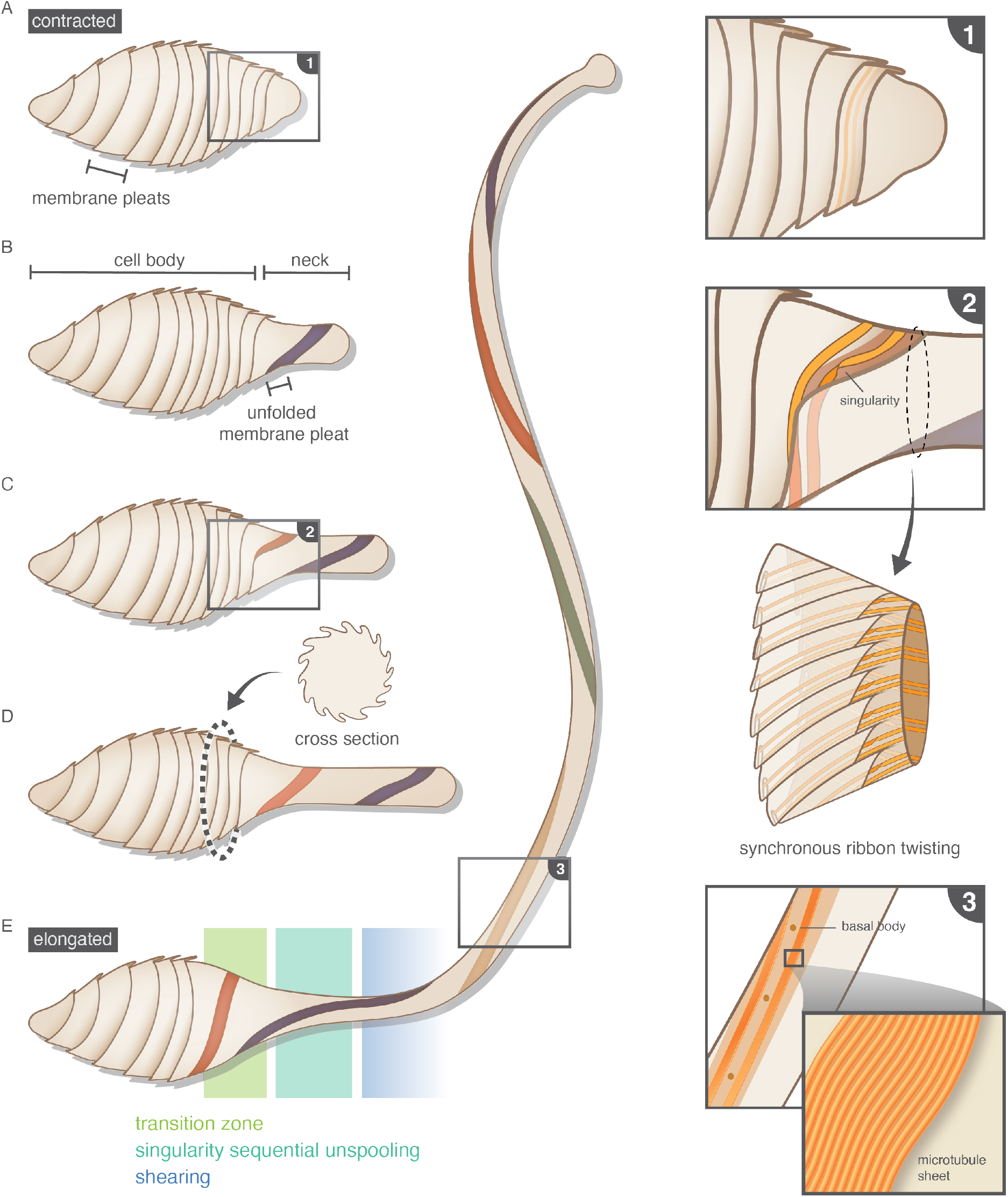
During extension, the microtubule filaments and the membrane pass through the transition zone for coordinated, reversible unspooling. This schematic shows a cell transitioning from a contracted to an elongated state. (A) In the first cell, the membrane is completely folded into pleats. Inset (1) from this schematic depicts microtubules stored in the pleats. (B) In the second cell, the neck is partially elongated, and one pleat is unfolded (shown in purple). In this cell, the transition zone between the cell body and the neck becomes apparent. (C) As the neck continues to elongate, more pleats unfold while in the transition zone. The specific location where the pleats transition from folded to unfolded occurs at point singularities, which is enlarged in inset (2). The synchronization of twist singularity during extension is shown below inset (2). (D) A cross section of the transition zone is shown to illustrate the 3-dimensional shape of the pleats. (E) In the final, most-elongated cell, we highlight the 3 regimes which are required for hyper-extension: the transition zone where the membrane is pleated, the zone where singularities propagate and the pleats sequentially unfold, and the last zone where the neck undergoes limited shear, resulting in thin hyper-extensions. In the neck of this last cell, inset (3) shows that the microtubule ribbons and the basal bodies are now most visible as the pleats are unfolded.

The work presented here also has implications in the design of macro, micro or nanorobotics. The stability and geometrical control of the barber pole architecture we described can be used for linear deployable structures, with applications ranging from space to medical robotics. It is remarkable to contemplate that *L. olor* undergoes repeated extreme morphological varia-tion tests (20k+ runs for a single cell), and is able to continually perform its desired function robustly. With our discovery of curved crease origami in a living cell, we intend to inspire similar robust, multi-stable abiotic structures where geometrical singularities encoded in origami patterns control unique kinetic behavior.

## Data Availability

All datasets are deposited at (DOI: 10.5061/dryad.4xgxd25g0).

## Acknowledgements

We thank all members of the Prakash lab (past and present) for helpful discussions. We particularly would like to thank Vishal Patil for scientific discussions on beam energetics, Scott Coyle and Deepak Krishnamurthy for general aspects of *L.olor* search behavior, and Hongquan Li for specialized imaging tools. We thank Rebecca Konte for assistance with and valuable feedback on figures. We thank the Berkeley Electron Microscopy Facility and specially Danielle for support during TEM data collection. We also thank staff members of CSIF Imaging facility at Stanford. E.F. acknowledges financial support from the NIH (T32GM008294-29). M.P. acknowledges financial support from NSF Career Award, Moore Foundation, HHMI Faculty Fellows program, NSF CCC (DBI-1548297) program, NSF Con-vergence Award (OCE-2049386), Schmidt Foundation, and CZ BioHub Investigators Pro-gram.

## Author Contributions

E.F and M.P designed the research. E.F. and M.P collected live imaging data. E.F collected fluorescence and TEM datasets and analyzed the same. E.F and M.P constructed origami models. E.F. and M.P wrote the manuscript.

