## Supplementary Information for "Curved crease origami and topological singularities at a cellular scale enable hyper-extensibility of *Lacrymaria olor*"

(Dated: August 3, 2023)

### CONTENTS

I. Tables 1

II. Methods 2
A. Cell Culture 2
B. DIC Imaging of Cells 2
C. Extension Rate and Strain Calculations 2
D. Filament Storage Calculation 2
E. Membrane Storage Calculation 3
F. Fluorescene Imaging 3
1. $\alpha$-tubulin Antibody Staining 3
2. Centrin Antibody Staining 4
3. Direct Membrane Labeling and Folds 4
G. Comparative Morphology of Cells with Protrusions: Isolation and Preparation 4
H. Radius and Helix Angle Measurements 4
I. TEM Imaging 4
J. 3D Projections 4
K. Energy Calculations 5
L. Cell Micromanipulation Assay 5
M. Paper Origami 5
N. Mylar and Bamboo Origami 6
O. Poisson Ratio Calculation 6

III. Supplementary Figures 6

IV. Supplementary Movies 13

References 14

### I. TABLES

| Cell | Diameter ($\mu$m) | Ext. Length ($\mu$m) | Ext. Time (s) | Max. Ext. Length ($\mu$m) |
|---|---|---|---|---|
| Fibroblast Cell (lamellipodium) [1] | 18 | 10 | 120 | 10 |
| Sea Urchin Embryonic Cell (filopodium) [2] | 5 | 80 | 480 | 80 |
| Dictyostelium (pseudopod) [3] | 12 | 5.4 | 11.6 | 5.4 |
| Neuron (axon) [4] | 15 | 64 | 86400 | 64 |
| Maize Gamete (pollen tube) [5] | 125 | 125000 | 37512 | 125000 |
| Nematocyte (nematocyst) [6] | 14.2 | 800 | 0.267 | 800 |
| Microsporidium (polar tube) [7] | 5 | 100 | 2 | 100 |
| Euglena [8] | 29 | 59 | 25200 | 59 |
| Tetrahymena (cilia) [9] | 35 | 6.5 | 7200 | 6.5 |
| Chlamydomonas (cilia) [9] | 10 | 10.7 | 14400 | 10.7 |
| Vorticella (stalk) [10] | 40 | 100 | 15 | 100 |
| Spirostomum [11] | 440 | 1100 | 1 | 1100 |
| Cultured Lacrymaria olor (neck) [12] | 40 | 100 | 1 | 700 |
| Wild-Type Lacrymaria olor (neck) [12–14] | 40 | 97.803 | 0.0652 | 1500 |

### II. METHODS

#### A. Cell Culture

*Lacrymaria olor* cell cultures were maintained and grown in the lab for use in experiments. The cells were obtained from Ryuji Yanase of Hyogo University, Japan [15]. Culture maintenance protocols were initially developed by Yanase et al, and further modified by Coyle et al [12, 15]. *L. olor* was grown in T225 TC-treated culture flasks (Corning). Each flask contained 200 mL of 0.01% Knop's at room temperature in the dark (Flinn Scientific). Cells were fed twice a week with 50 mL of concentrated Cyclidium cultures per flask.

Cyclidium cultures were also grown in the lab. Cyclidium were initially grown in 300 mL flasks containing a nutritional agar pad (2% Agar, 1 mg/ml Yeast Extract, 1 mg/ml Skim Milk) and 250 mL 0.01% Knop's at room temperature in the dark. 3 days prior to feeding, 2 Cyclidium flasks were transferred to 8x160mm Petri dishes, still maintained at room temperature in the dark. The day of feeding, the Cyclidium were removed from the Petri dishes, and were concentrated and cleaned by passing the cells through a 70 $\mu$m filter, and then through a 40 $\mu$m filter. After each filtering, the Cyclidium cells were spun down at 500xRCF, and then were washed twice with fresh 0.01% Knop's. With the final concentrated and cleaned Cyclidium, new cultures were started using a 1:100 ratio of concentrated culture : fresh 0.01% Knop's in the 300 mL flasks. The remainder was put into the T225 flasks containing *Lacrymaria olor.*

*Lacrymaria olor* cells were removed for experiments from their cultures after feeding. 25-50 mL of culture was removed and concentrated at 500xRCF for 5 minutes. In 24 hours, the cells were then ready for imaging. The time after feeding was required as the necks were often damaged during the concentration.

#### B. DIC Imaging of Cells

Cells were placed in chambers of height 50 $\mu$m and were imaged using DIC and a high-speed camera (200 fps).

#### C. Extension Rate and Strain Calculations

Cell types were chosen for comparison to *L. olor* if they had uniaxial deformations. Extension rates were obtained in the literature for all cell types except for *L. olor.* Lamellipodium measurements were obtained on NIH 3T3 fibroblast cells in mice [1]. Filopodia measurements were taken on sea urchin embryos [2]. Dictyostelium measurements were taken from the "fruiting body" growth stage[3]. Neuron measured from mouse adult C57B16/J animals [4]. Polar tube measurements were obtained in maize grains [5]. Nematocyst measurements were obtained from nematocytes [6]. Microsporidium (*Nosema algerae*) growth rates were obtained from ejection of their polar tubes [7]. *Euglena gracilia* whole cell morphodynamic rates were used [8]. Cilia growth rates were obtained for both *Tetrahymena* and *Chlamydomonas* [9]. Vorticella measurements are from post-contraction stalk extensions [10]. Spirostomum measurements are for whole cell relaxation post contraction [11]. Strain measurements were calculated for all cell types by computing $\Delta$L/L, where $\Delta$L is the length of the deformation, and L is the diameter of the cell body prior to deformation. Extension rate and strain for cultured *L. olor* were measured using videos of live cells obtained in the lab. Wild-type *L. olor* measurements were made using Youtube videos posted online on several public accounts [16–21]. Scales were created from these videos assuming that the cell body diameter is 40 $\mu$m, since this value is highly consistent within our own data of *L. olor.*

#### D. Filament Storage Calculation

We approximate the cell body as a cylinder with both a diameter and a height of 40 $\mu$m (Fig. S4A):
The surface of such a cylinder is represented by

$$\mathrm{SA_c} = 2\pi r_c h_c$$

where $r_c$ is the radius of 20$\mu$m, and $h_c$ is the 40$\mu$m height of the cylinder. If a single microtubule filament of thickness 25 nm is wrapped around this cylinder (Fig. S4B), the surface area of this single filament is represented by

$$\mathrm{SA_f} = 2\pi r_\mathrm{c} t_\mathrm{f}$$

If this filament is maximally packed on the cylinder, the number of wraps this filament can make is the ratio of the surface area of the cylinder over the surface area of a single filament:

$$n_\mathrm{wraps} = \frac{2\pi r_\mathrm{c} h_\mathrm{c}}{2\pi r_\mathrm{c} t_\mathrm{f}} = \frac{h_\mathrm{c}}{t_\mathrm{f}} = 1600$$

The length of a microtubule which wrap 1600 times around a cylinder of radius 20$\mu$m and height 40 $\mu$m is then:

$$L_\mathrm{fil} = n_\mathrm{wraps}(2\pi r) = 201,061\mu\mathrm{m}$$

In the literature [13, 14], and from our own data, we know that instead of just a single filament wrapping around the cell, there are 15 microtubule "filaments." Adjusting our above calculation to account for 15 filaments (Fig. S4C), the number of wraps is now represented by:

$$n_\mathrm{wraps} = \frac{2\pi r_\mathrm{c} h_\mathrm{c}}{15(2\pi r_\mathrm{c} t_\mathrm{f})} = 106$$

The length of a single filament which can then be stored in this cylinder is 13,404 $\mu$m.

If we update this approximation again considering that these 15 filaments are actually 15 pairs of filaments, which we observed in our fluorescence data, the number of wraps is now:

$$n_\mathrm{wraps} = \frac{2\pi r_\mathrm{c} h_\mathrm{c}}{2 * 15 * (2\pi r_\mathrm{c} t_\mathrm{f})} = 53$$

giving a maximum stored filament length of 6,702 $\mu$m (Fig. S4D).

Our TEM data allows us to make yet another adjustment to this calculation. We observed that the filaments are actually sheets of 14 microtubules. This requires us to adjust the surface area for a single filament (Fig. S4E):

$$\mathrm{SA_f} = 2\pi r_\mathrm{c}(14 t_\mathrm{f})$$

Our number of wraps is then 3.8 per pair of of a sheet of filaments, giving a maximum deployable length of 478 $\mu$m.

Our last adjustment which we can make is accounting for the membrane folds which layer the microtubule filaments. We observed up to 3 layers of paired filament sheets inside the cell, increasing the total number of wraps 3-fold, and allowing a maximum deployable length of 1436 $\mu$m (Fig. S4F-G). This approximation is the same order of magnitude as the maximum observed neck length in *L. olor*.

### E. Membrane Storage Calculation

Assuming the neck is a 1500$\mu$m-long cylinder with a 5$\mu$m diameter, the surface area of the neck is about 23,000 $\mu$m$^2$. Assuming the cell body is a sphere with a 40$\mu$m diameter, the cell body would have a much smaller surface area of about 5000 $\mu$m$^2$.

### F. Fluorescene Imaging

#### 1. *$\alpha$-tubulin Antibody Staining*

Cells were isolated in 96-well plates, and then fixed for 30 minutes in 100 $\mu$L of fixative solution (2% (w/v) PFA and 0.5% (w/v) Triton X-100 in PHEM buffer). The fixative solution was then replaced with 0.05 M glycine in PBS for 20 minutes. The glycine solution was then replaced with blocking solution (0.2% bovine serum albumin (w/v) in PBS) for 30 minutes. Blocking solution was then replaced by antibody solution (0.1% (w/v) BSA, 0.1%(w/v) Triton X-100 in PBS), which also contained the Alexa Fluor 488 conjugated anti-alpha-tubulin antibody (Invitrogen, RRID: AB2532182) diluted 1:500, for 1 hour. Cells were then washed 3 times in PBS containing 0.2%(w/v) Triton X-100. Cells were lastly mounted onto glass slides. Images were collected using an LSM 780 microscope. .

##### 2. Centrin Antibody Staining

We stained fixed cells with anti-centrin mouse monoclonal antibody as a primary antibody (Milipore, RRID: AB10563501) and Alexa Fluor 568 conjugated anti-mouse IgG antibody (Invitrogen, RRID: AB2534072) as a secondary antibody using the same protocol for the microtubule staining and imaged again on a LSM780 Confocal Microscope.

##### 3. Direct Membrane Labeling and Folds

We labeled fixed cells with the membrane stain FM-464 (Thermofisher, T13320).

#### G. Comparative Morphology of Cells with Protrusions: Isolation and Preparation

*Dileptus sp.* and *Tracheloraphis sp.* were both collected and isolated in from local freshwater sources. They were then fixed and stained according to the microtubule staining section above.

#### H. Radius and Helix Angle Measurements

The radius and helix angle were measured from the $\alpha$-tubulin fluorescence images. To compute the radius, we used a curve tracing the outline of the cell as well as a curve tracing the center of the cell from tail to oral apparatus. The radius is the distance from each point on the curve tracing the outline of the cell to its closest centerline point. To compute the helix angle, we chose several arbitrary points on each visible filament, and measured the angle between the tangent to the filament at that point and the tangent to the centerline at that point.

#### I. TEM Imaging

Cells were isolated in a 96-well plate, and then prepared for TEM using a protocol developed by the Berkeley Electron Microscopy Lab. Cells were fixed with an EM-grade fixative solution (2% PFA, 2% Glutaraldehyde in 0.1% Knop's) for 1 hour at RT (Electron Microscopy Sciences). A drop of SafroninO was added to the solution containing the cells to dye the cells so they are visible. The cells were then moved to a glass dish with shallow wells sitting on ice, and any excess solution was removed. 2% melted agarose was then added to the shallow well containing the cell, and was mixed. Any air bubbles were removed. Once the gel solidified, each cell was cut out using a scalpel, and were placed in eppendorf tubes containing 2% glugaraldehyde and 2% PFA in PBS. The samples were then rinsed in PBS 3 times for 5 minutes each on a shaker. The PBS was then replaced with PBS containing 1% Osmium tetroxide and 1.6% Potassium ferricyanide (KFECn). The cells were again rinsed 3 times in PBS, and one time in distilled water. Cells were then dehydrated in increasing increments of acetone (35%, 50%, 70%, 80%, 95%, 100%, 100%) for 10 minutes each. The cells were then infiltrated in 2:1 acetone:resin containing the accelerator BDMA for 1 hour. This solution was then replaced with a 1:1 acetone:resin containing BDMA for 1 hour, and again with 1:2 acetone:resin containing accelerator overnight. Then the cells were incubated in 100% resin containing BDMA 3 times for 1-2 hours each. Lastly, the cells were embedded into a mold, and baked in the oven at 60 for 2 days. Cells were then sectioned into 70 $\mu$m slices, and were imaged using a FEI Tecnai 12 Transmission Electron Microscope.

#### J. 3D Projections

3D projections were made using confocal microscopy z-stack images of fixed cells stained with an $\alpha$-tubulin conjugated antibody. The z-stacks were max-projected into a single 2d-image. Filaments were then traced by hand using an iPad and Apple Pencil. The outline of the cell on either side, and an approximate centerline along the length of the cell were traced as well using this method. We assume that the body is axisymmetric down the centerline, which allows us to set the z-value of any point on the centerline as the distance from that point in x,y to the closest point on the traced outline of the cell, where the z-value will remain 0. For each point on each filament, we then compute a z-value by forming a circle in 3d from the closest point along the outline of the cell and the closest point on the centerline, and finding the closest value on this circle in x,y to the point on the filament. The z-value of the point on the filament is then assigned the z-value of the circle at that point.

#### K. Energy Calculations

To understand how *L. olor* is able to achieve such rapid, large deformations, we compute the bending and twisting energy of the microtubule bundles, using discrete forms of the following equations:

$$E_{\text{bend}} = \int_0^L (EI_x k_1^2 + EI_y k_2^2)\, ds$$

$$E_{\text{twist}} = \int_0^L GJk_3^2\, ds$$

where $E$ is the bending modulus of microtubule filaments [22], G is the shear modulus [23], $I_x$ and $I_y$ are the moment of inertia of the bundle in x and y, respectively, $J$ is the area moment of inertia, and $k_1$, $k_2$, and $k_3$ come from the bending and twist of the material frame along the length of the cell. We assume that both microtubule filament and total membrane are conserved, and that the microtubule bundles remain in the same plane as the membrane, or that the filaments do not twist with respect to the membrane such that any microtubule twist is paired with twist of the membrane as well. The moment of inertia is computed assuming there are 14 filaments in a microtubule bundle. We calculated this number using measurements taken on the TEM data. The medians for the length of the microtubule bundles in contracted and elongated cells were 0.386 $\mu$m and 0.408 $\mu$m, respectively. In these images, we also measured the diameter of an individual filament as 16.61 nm +/- 3.115 nm, and the distance of the gap between the filaments as 11.46nm +/- 3.081 nm. To compute the number of filaments in each bundle, we divide 386 nm and 408 nm by the sum of our diameter and gap distance (16.61 nm + 11.46 nm = 28.07 nm), which is 28.07 nm, giving a solution of 13.75 and 14.53 filaments. We therefore use a bundle of a rounded value of 14 filaments to compute the moments of inertia. We then define a local basis $\{d_1, d_2, d_3\}$, where $d_3$ is the unit tangent vector of the bundle, $d_1$ is the unit surface normal, and $d_2$ is the third vector in this orthonormal basis. We use this local material frame, and assuming inextensibility of our discrete filament, compute the $k_i$ values, and to compute the bending and twist energies.

#### L. Cell Micromanipulation Assay

We performed micromanipulation stretching assays of *L. olor* to observe membrane and shape dynamics during extension. We held the cell using suction pressure (Sutter Xenoworks Microinjector) with two holding micropipettes (Sutter P-97 Micropipette Puller). While maintaining one micropipette in a fixed position, we began moving the second micropipette away from the cell using a micromanipulator (Scientifica PatchStar Micromanipulator, Narishige MMO-203 Three-axis Oil Hydraulic Micromaniuplator). We were able to increase the neck length of cells up to 200+ $\mu$ms in several versions of this experiment. We observed that as we elongate the cell neck, the cell body and neck rotate in response. We measured this rotation, observing that for a neck extension of 47.45 $\mu$m, from 126.23 to 173.68 $\mu$m in 4.063s, the cell rotates 128.28, rotating at a speed of 31.57/s (Supplementary Video 4).

#### M. Paper Origami

Origami cylinders were made using 25 cm x 25 cm sheets of origami paper. Patterns traced by pen to create creases on both sides of the paper. The pleats were colored with marker to highlight the membrane which is accessible post elongation. During extension, one side was kept in the open configuration to seed d-cones and therefore extension at that end. Force was generated using strings and by pulling manually.

#### N. Mylar and Bamboo Origami

Origami cylinders were made using 12in x 12in mylar sheets. Patterns traced by pen to create creases on both sides of the paper. Bamboo was taped to the surface to represent filaments. During extension, one side was kept in the open configuration to seed d-cones and therefore extension at that end. Force was generated using strings and manual pulling.

#### O. Poisson Ratio Calculation

We computed the poisson ratio of the origami cylinders by measuring the diameter ($d_i$,$d_f$) and length ($l_i$,$l_f$) of cylinders in the fully folded state and the fully unfolded state. We then compute the poisson ratio using the following expression:

$$v = -(d_f - d_i/d_i)/(l_f - l_i/l_i)$$

### III. SUPPLEMENTARY FIGURES

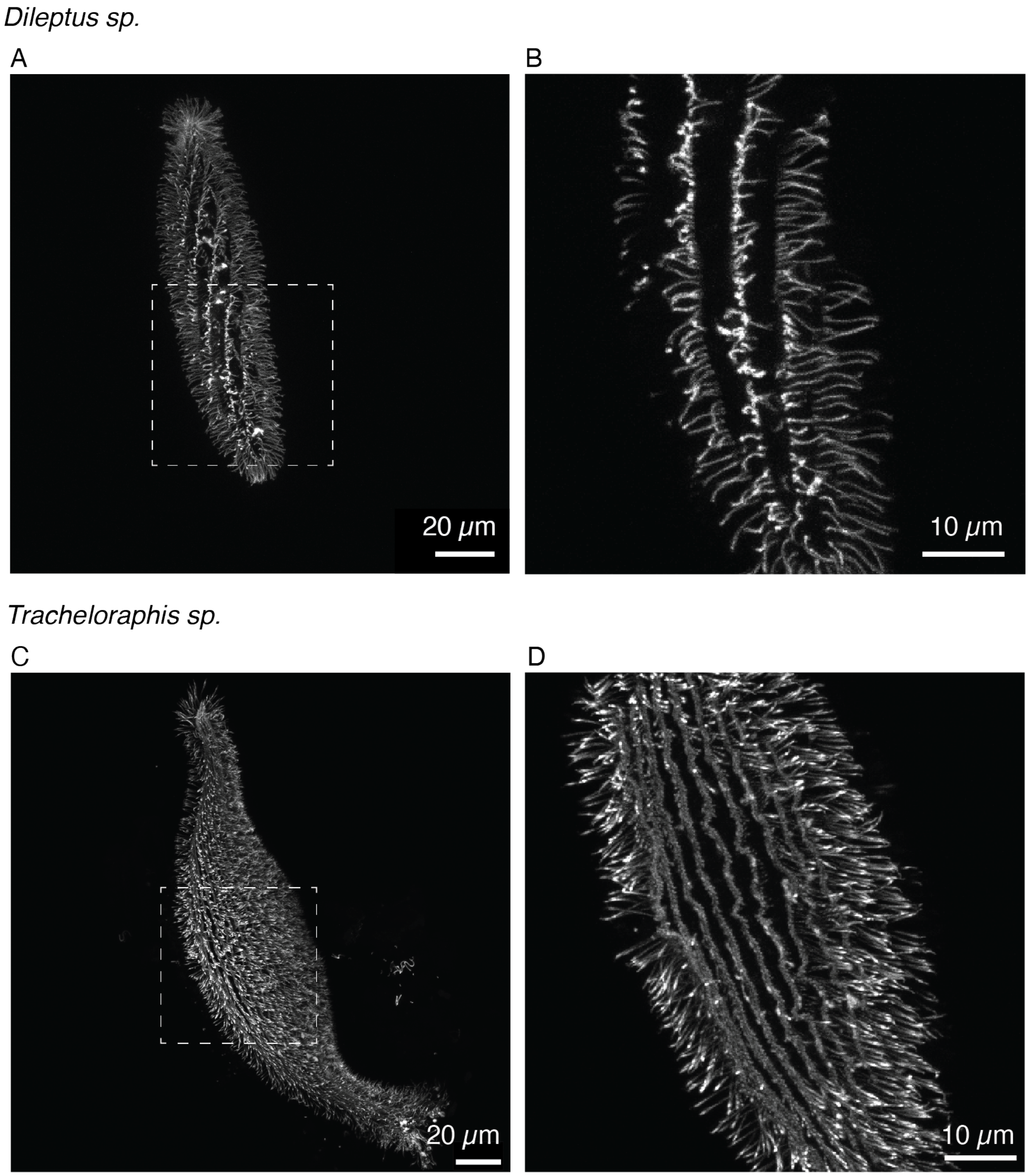


**FIG. S1 The cortical cytoskeleton of ciliates which have similar cell morphologies but non-extensile morphology dynamics are uniform along the length of the cell.** Z-projections of fixed *Dileptus sp.* and *Tracheloraphis sp.* stained with $\alpha$-tubulin. (a) Whole cell image of *Dileptus sp.*/ (b) Enlargement of boxed region from a single z-slice of (a). (c) Whole cell image of *Tracheloraphis sp.*. (d) Enlargment of boxed region from a single z-slice of (c). are whole cell images, and the right are enlargements of the boxed regions. In both cells the microtubule filaments align along the length of the cell.

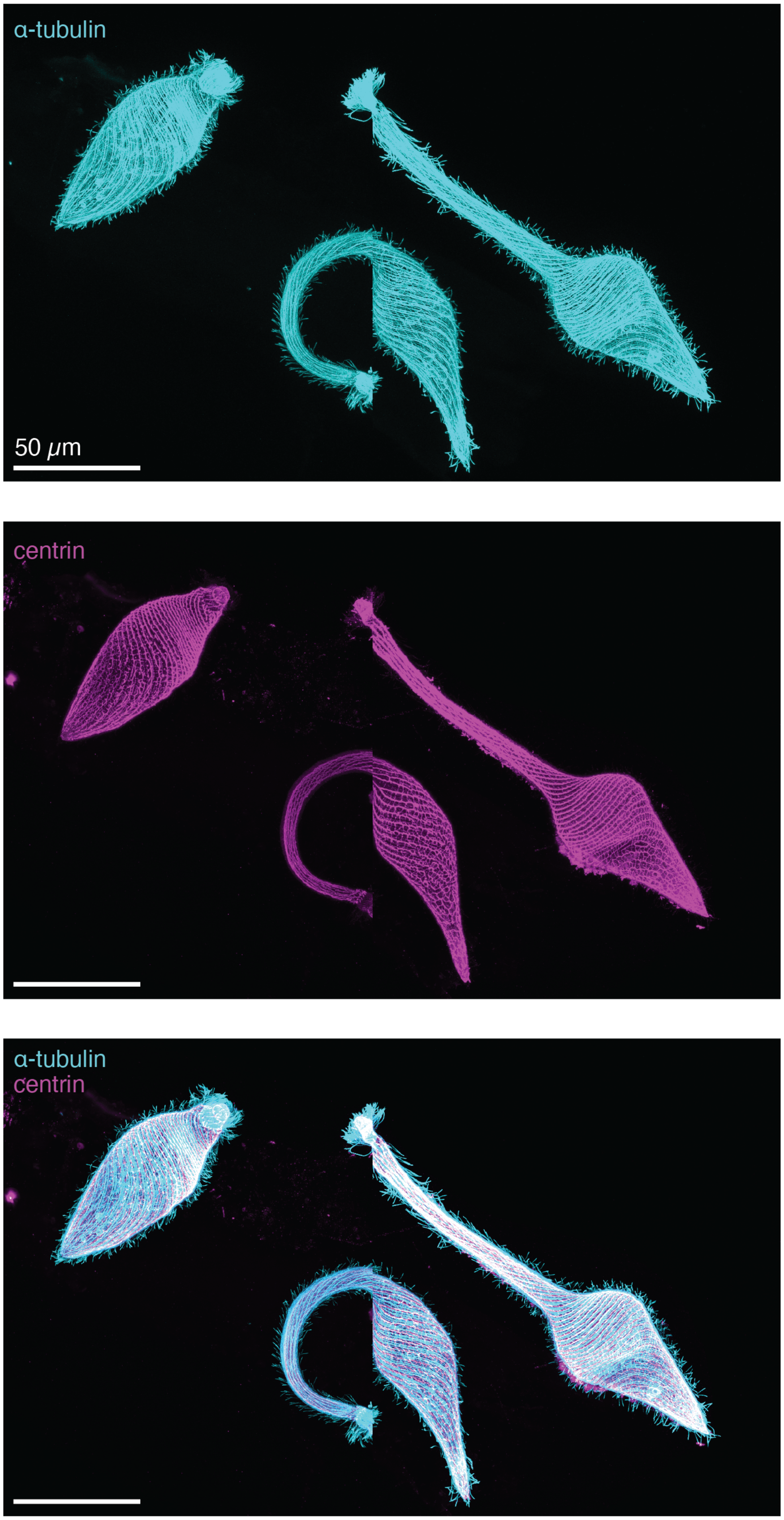


**FIG. S2 The cortical cytoskeleton Lacrymaria olor contains $\alpha$-tubulin and centrin, both arranged in a helical architecture.** Z-projections of fixed *L. olor* stained with $\alpha$-tubulin and centrin antibodies.

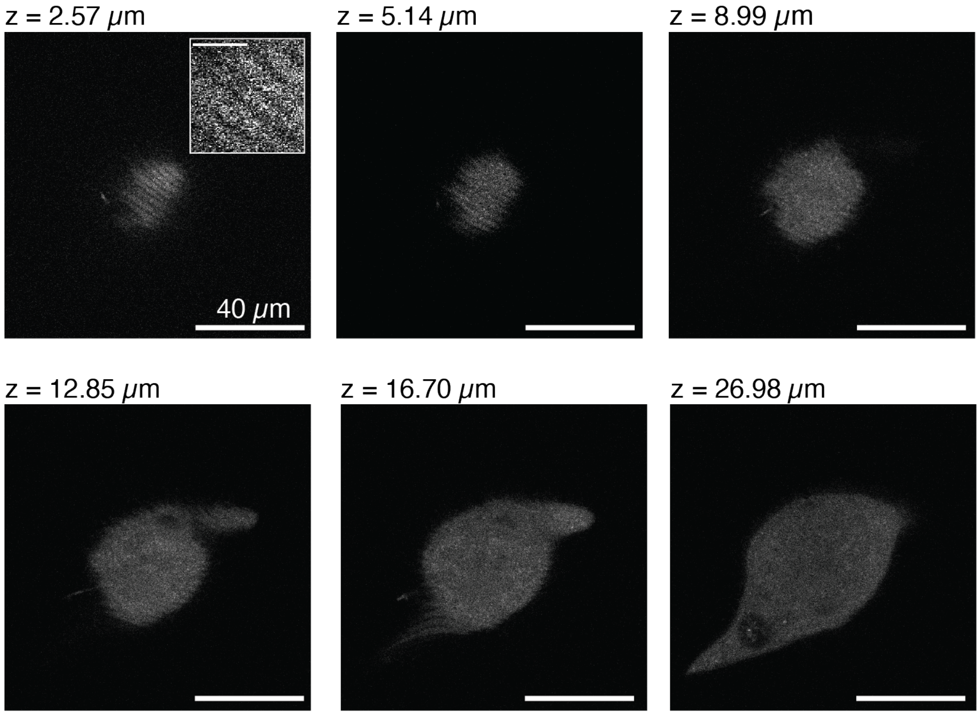


**FIG. S3 Membrane stains reveal membrane folds which are present in the cell body.** Confocal z-stack of FM4-64 Stain, with z distance as the distance from the surface of the cell. Membrane folds are visible in the top 3 panels.

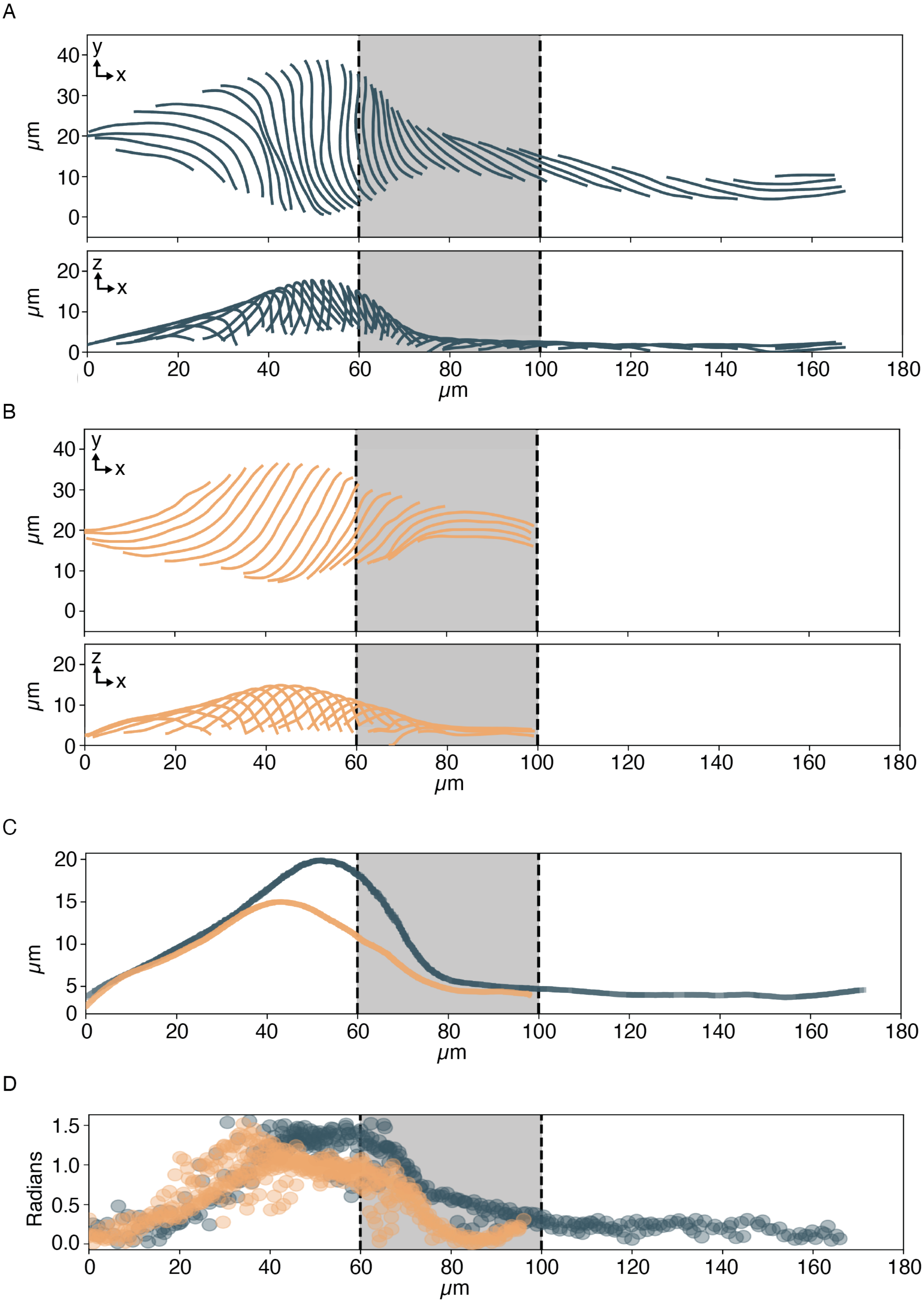


**FIG. S4** **Filament traces are projected into 3D assuming axisymmetry.** (a) The filament traces and their 3d shape for an elongated cell. (b) The filament traces and their 3d shape for a contracted cell. (c) The radius for each 3d-projected point in each filament. (d) The helix angle for each 3d-projected point in each filament.

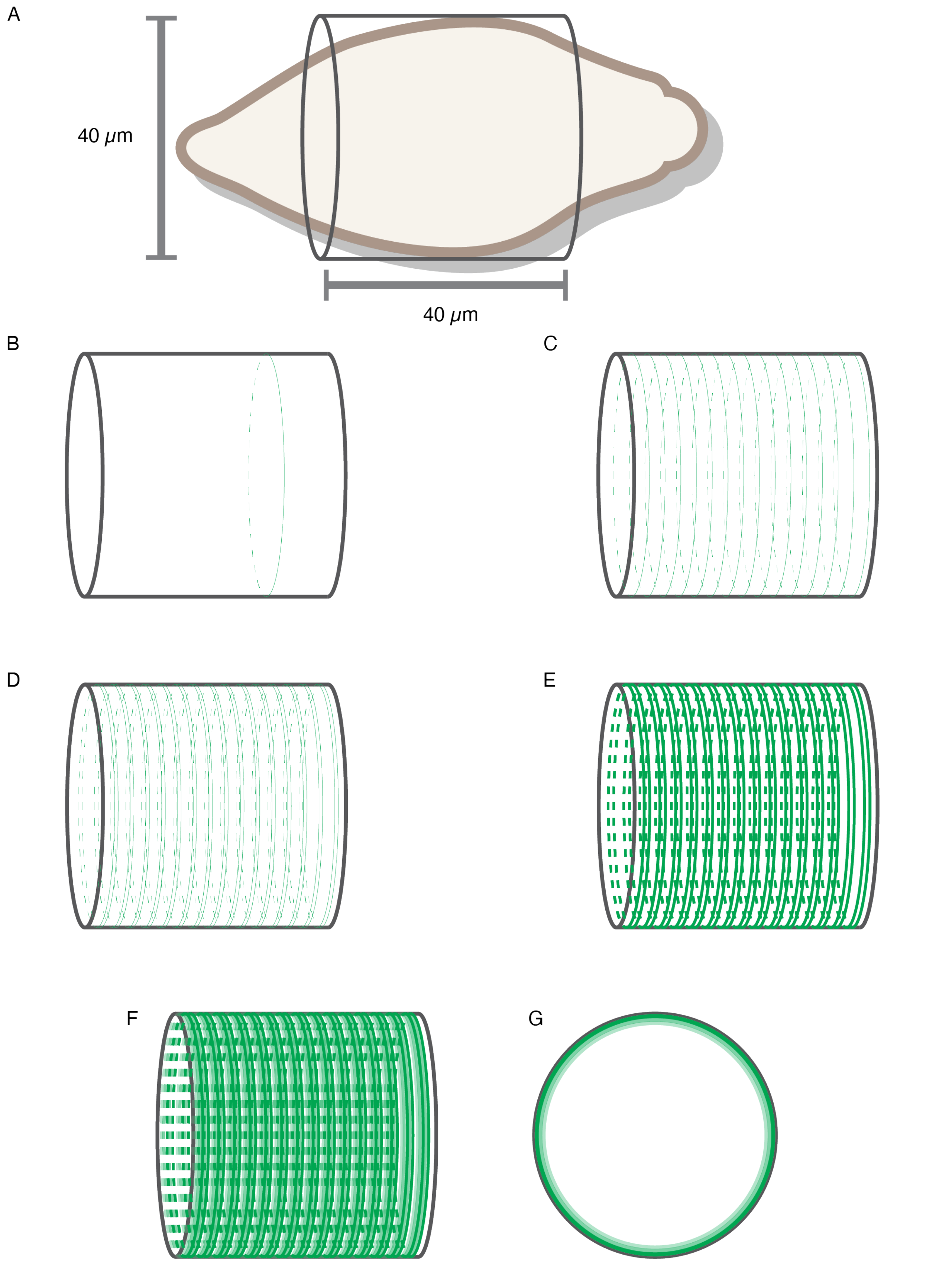


**FIG. S5 Microtubule storage approximation in a single cell.** (a) We approximate a contracted cell as a cylinder. (b) A schematic of a single filament wrapped around the cylinder. (c) A schematic of 15 filaments wrapped around the cylinder. (d) A schematic of 15 pairs of filaments wrapped around the cylinder. (e) A schematic of 15 pairs of microtubule sheets wrapped around the cylinder. (f) A schematic of 15 pairs of layered microtubule sheets wrapped around the cylinder. (g) A cross-section view of a one pair of layered microtubule sheets.

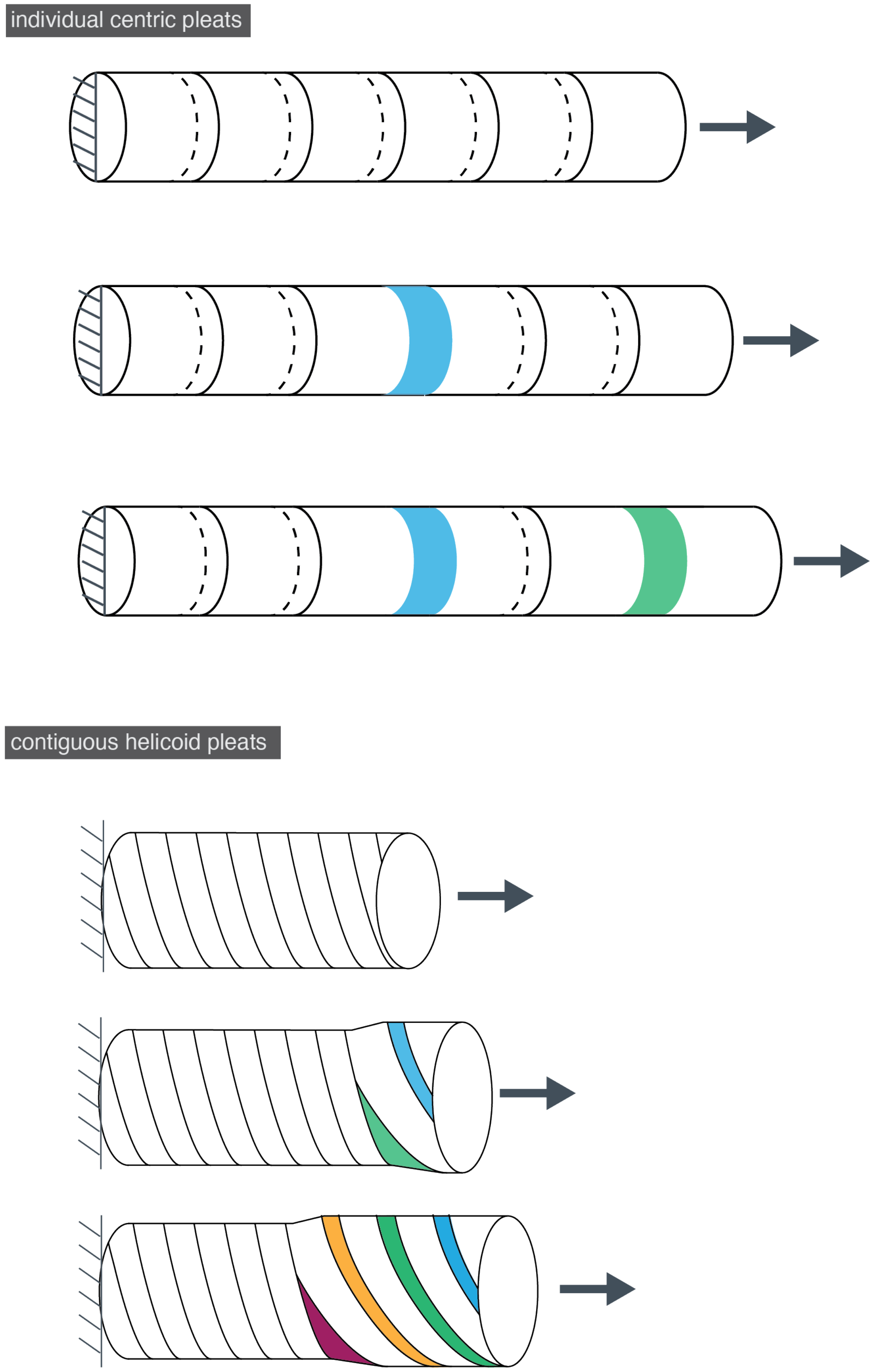


**FIG. S6** **Helicoid contiguous pleats enable sequential unspooling whereas indivual centric pleats stochastically unfold.**

### IV. SUPPLEMENTARY MOVIES

**Supplementary Movie 1:** High speed cellular extension from contracted to elongated states
**Supplementary Movie 2:** Microtubule layering along the length of the cell
**Supplementary Movie 3:** Micromanipulation-induced neck extensions
**Supplementary Movie 4:** Sequential unspooling of curved crease origami

[1] Milo, R., Jorgensen, P., Moran, U., Weber, G. & Springer, M. Bionumbers–the database of key numbers in molecular and cell biology. *Nucleic acids research* **38**, D750–3 (2010).

[2] Svitkina, T. M. *Filopodia and Lamellipodia*, vol. 2, 683–693 (Elsevier Inc., 2016).

[3] van Haastert, P. J. Unified control of amoeboid pseudopod extension in multiple organisms by branched f-actin in the front and parallel f-actin/myosin in the cortex. *PLoS ONE* **15** (2020).

[4] Santos, T. E. *et al.* Axon growth of cns neurons in three dimensions is amoeboid and independent of adhesions. *Cell Reports* **32** (2020).

[5] House, L. R. & Nelson, O. E. Tracer study of pollen-tube growth in cross-sterile maize. URL https://academic.oup.com/jhered/article/49/1/18/801280.

[6] Colin, S. P. & Costello, J. H. Functional characteristics of nematocysts found on the scyphomedusa cyanea capillata. *Journal of Experimental Marine Biology and Ecology* **351**, 114–120 (2007).

[7] Frixione, E. *et al.* Dynamics of polar filament discharge and sporoplasm expulsion by microsporidian spores. *Cell Motility and the Cytoskeleton* **22**, 38–50 (1992).

[8] Lonergan, T. A. Regulation of cell shape in euglena gracilis i. involvement of the biological clock, respiration, photosynthesis, and cytoskeleton the involvement of respiratory and photosynthetic pathways in the celi shape changes was investigated with energy pathway inhibitors. antimycin (1983). URL https://academic.oup.com/plphys/article/71/4/719/6078962.

[9] Reynolds, M. J. *et al.* The developmental process of the growing motile ciliary tip region. *Scientific Reports* **8** (2018).

[10] Ryu, S., Pepper, R. E., Nagai, M. & France, D. C. Vorticella: A protozoan for bio-inspired engineering (2017).

[11] Mathijssen, A. J., Culver, J., Bhamla, M. S. & Prakash, M. Collective intercellular communication through ultra-fast hydrodynamic trigger waves. *Nature* **571**, 560–564 (2019).

[12] Coyle, S. M. Ciliate behavior: blueprints for dynamic cell biology and microscale robotics. *Molecular Biology of the Cell* **31**, 2415–2420 (2020).

[13] Müller, O. Zoologicae danicae prodromus, seu animalium daniae et norvegiae indigenarum characteres, nomina et synonyma imprimis popularium. *Havniae* **32**, 1–282 (1776).

[14] Müller, O. F. & Fabricius, O. *Animalcula infusoria fluviatilia et marina que detexit, systematice descripsit et ad vivum delineari curavit Otho Fridericus Müller sistit opus hoc posthumum quod cum tabulis Aeneis L. in lucem tradit vidua ejus nobilissima cura Othonis Fabricii* (Typis N. Mölleri, 1786).

[15] Yanase, R., Nishigami, Y., Ichikawa, M., Yoshihisa, T. & Sonobe, S. The neck deformation of lacrymaria olor depending upon cell states. *Journal of Protistology* (2018).
